# Hemicentin mediated type IV collagen assembly strengthens juxtaposed basement membrane linkage

**DOI:** 10.1101/2021.12.21.473673

**Authors:** Claire A. Gianakas, Daniel P. Keeley, William Ramos-Lewis, Kieop Park, Ranjay Jayadev, Qiuyi Chi, David R. Sherwood

## Abstract

Basement membrane (BM) matrices surround and separate most tissues. However, through poorly understood mechanisms, BMs of adjacent tissues can also stably link to support organ structure and function. Using endogenous knock-in fluorescent proteins, conditional RNAi, optogenetics, and quantitative live imaging, we identified matrix proteins mediating a BM linkage (B-LINK) between the uterine utse and epidermal seam cell BMs in *Caenorhabditis elegans* that supports the uterus during egg-laying. We found that hemicentin is secreted by the utse and promotes fibulin-1 assembly to jointly initiate the B-LINK. During egg-laying, however, both proteins decline in levels and are not required for B-LINK maintenance. Instead, we discovered that hemicentin also promotes type IV collagen assembly, which accumulates to high levels during egg-laying and sustains the B-LINK during the mechanically active egg-laying period. This work reveals mechanisms underlying BM-BM connection maturation and identifies a crucial function for hemicentin and fibulin-1 in initiating attachment and type IV collagen in strengthening this specialized form of tissue linkage.

**Summary:** Tissue attachment through linking juxtaposed basement membranes (BMs) is crucial for the structure and function of many organs. Gianakas et al. identify a key role for hemicentin and fibulin-1 in initiating BM-BM attachment and type IV collagen in stabilizing this linkage and allowing it to resist high mechanical loads.

## Introduction

Basement membranes (BMs) are thin dense sheets of extracellular matrix that surround most tissues and provide structural and signaling support (Jayadev and Sherwood, 2017). Two of the most abundant core BM components are laminin and type IV collagen, which form independent self-oligomerizing networks. Laminin initiates BM formation and associates with cells through binding to integrin and dystroglycan receptors (Li et al., 2017). Type IV collagen networks have high tensile strength and allow BMs to mechanically support tissues (Fidler et al., 2018). BMs also contain other core components such as perlecan, agrin and nidogen, which connect the collagen and laminin networks (Hohenester and Yurchenco, 2013), and harbor matricellular proteins, proteases, and growth factors (Glentis et al., 2014; Keeley et al., 2020; Pozzi et al., 2017). Although BMs usually separate tissues, in some instances, BMs of adjacent tissues stably link and connect tissues to structurally support organs and facilitate specialized functions. One of the best characterized instances occurs in the kidney, where epithelial podocytes and the vascular endothelium synthesize distinct BMs that fuse to form the glomerular BM, which filters blood (Abrahamson, 1985; Keeley and Sherwood, 2019). Stable BM-BM linkages occur between other tissues as well (Keeley and Sherwood, 2019), including the alveolar and capillary BMs in the lung that facilitate gas exchange (Vaccaro and Brody, 1981; Makanya et al., 2013), and the capillary endothelial and parenchymal BMs of brain astrocytes that help form the blood-brain barrier (Sixt et al., 2001; Obermeier et al., 2013). Despite the importance of BM-BM attachments in stably joining tissues, little is known about how these connections are made and maintained because of the complexity of BMs, the difficulty in examining tissue interactions, and the lack of in vivo models that allow visualization and perturbation of BM components.

*C. elegans* is a powerful experimental model to understand how tissues attach through BM-BM linkages. *C. elegans* are optically transparent, have simple tissues, and BM components can be selectively targeted through conditional RNAi knockdown to determine time of function. Furthermore, nearly all BM components have been endogenously tagged with genetically encoded fluorophores, which allows for visualization and quantitative analysis of localization, levels, and dynamics (Keeley et al., 2020). We previously characterized a small, transient BM-BM linkage between the gonadal and vulval BMs beneath the *C. elegans* anchor cell, which we termed a B-LINK (for Basement membrane LINKage) (Morrissey et al., 2014). B-LINK formation allows the anchor cell to invade through and degrade both BMs simultaneously to initiate uterine-vulval tissue connection (Sherwood and Sternberg, 2003; Hagedorn et al., 2013). The extracellular matrix protein hemicentin (*C. elegans* HIM-4) is secreted by the anchor cell just prior to invasion and forms punctae that reside between the adjacent BMs to promote their connection (Morrissey et al., 2014). The integrin heterodimer (*α*INA-1/*β*PAT-3) organizes hemicentin into punctae and the plakin cytolinker (VAB-10A) tethers the anchor cell to hemicentin (Morrissey et al., 2014). Suggesting a common role in BM-BM linkage, zebrafish hemicentins facilitate transient BM-BM connections during development between epithelia in fin epidermis and between somites and epidermis (Carney et al., 2010; Feitosa et al., 2012).

Most BM-BM linkages in vertebrates are stable, present in mechanically active tissues, and subject to stresses from blood flow and muscle contractions (Keeley and Sherwood, 2019). It is unclear whether these long-term BM-BM linkages require additional matrix components to stabilize the connection. In addition to the anchor cell B-LINK, we identified a long-term B-LINK between the *C. elegans* uterine utse cell and epidermal seam cells that links the uterus to the epidermis for the lifetime of the animal (Morrissey et al., 2014). The utse-seam B-LINK is thought to help the uterus withstand the mechanical stresses of egg-laying (Newman et al., 1996; Schindler and Sherwood, 2013). All anchor cell B-LINK components localize to the utse-seam B-LINK, and their loss results in the uterine tissue and other internal organs expelling from the animal after initiation of egg-laying, suggesting they are also required for this long-term BM-BM linkage (Morrissey et al., 2014; Vogel and Hedgecock, 2001). The matricellular protein fibulin-1 (FBL-1) also localizes to the utse-seam B-LINK, but its role in BM-BM linkage has not been examined (Muriel et al., 2005). Whether additional BM matrix components might contribute to this long-term B-LINK that supports a mechanically active tissue is not known.

Using endogenous fluorophore knock-in strains for hemicentin, α-integrin, and plakin, we found that the utse-seam B-LINK forms during the L4 larval stage, just prior to adulthood and egg-laying. Optogenetic inducible muscle contraction and RNAi-mediated knockdown revealed that hemicentin is required for the uterus to resist the physical stress of muscle contraction and egg-laying, providing direct evidence that the B-LINK anchors the uterus during egg passage. Through visual and RNAi screening, we discovered that the matrix proteins fibulin-1, perlecan (UNC-52) and type IV collagen (EMB-9), are also components of the utse-seam B-LINK. Conditional and tissue-specific RNAi knockdown revealed that hemicentin is secreted by the utse and promotes the assembly of fibulin-1, type IV collagen and perlecan. Quantitative imaging revealed that fibulin-1 and hemicentin reach their peak levels at the B-LINK at the onset of egg-laying, and loss of fibulin-1 and hemicentin caused early defects in utse-seam cell attachment, while later RNAi targeting did not affect the B-LINK. In contrast, type IV collagen and perlecan reached maximal levels later, during the peak of egg-laying, and type IV collagen loss first perturbed the B-LINK after egg-laying was underway, suggesting that it helps resist the high mechanical stress of egg-laying. These findings identify matrix components required for a stable BM-BM linkage that resists mechanical stress and uncover distinct roles in mediating the initial formation and maintenance of a stable BM-BM linkage.

## Results

### The utse and seam cells are connected by a B-LINK during the L4 larval stage

The utse-seam BM-BM interface forms a BM-BM linkage, which we have termed a B-LINK (Fig. 1, A-C) (Morrissey et al., 2014; Vogel and Hedgecock, 2001). A key component of the B-LINK is hemicentin (*C. elegans* HIM-4), a large (5198-residue) conserved matrix protein, that localizes to the utse-seam interface (Fig. 1 C, Movie S1). The utse-seam B-LINK is also composed of the transmembrane integrin heterodimeric receptor (αINA-1/βPAT-3), and the cytosolic cytolinker plakin (VAB-10A) (Morrissey et al., 2014) (Fig. 1 C) and is thought to maintain the uterus within the body cavity and prevent uterine prolapse during egg-laying. The utse cell forms from fusion of ventral uterine cells that contact the seam cells during the early L4 larval stage (Ghosh and Sternberg, 2014). However, it is not known when the B-LINK first develops to connect these two tissues. To determine when the utse-seam B-LINK forms and how it develops over time, we examined the region where the utse and seam cells come into contact (visualized with *cdh- 3p::mCh::PH* and *scmp::GFP::CAAX*, respectively). We found the utse and seam are already in contact by the mid-L4 stage, at which point the length of contact measures ∼50 µm and the utse midsection length is ∼25 µm (Fig. 2 A and B). The length of utse-seam contact and the utse midsection continued to extend in the late-L4 stage and by the young adult the utse-seam contact was ∼60 µm in length and the midsection was ∼45 µm long. Additionally, by the young adult stage the end of each utse arm turned inwards away from the seam (Fig. 2 B).

**Figure 1.**
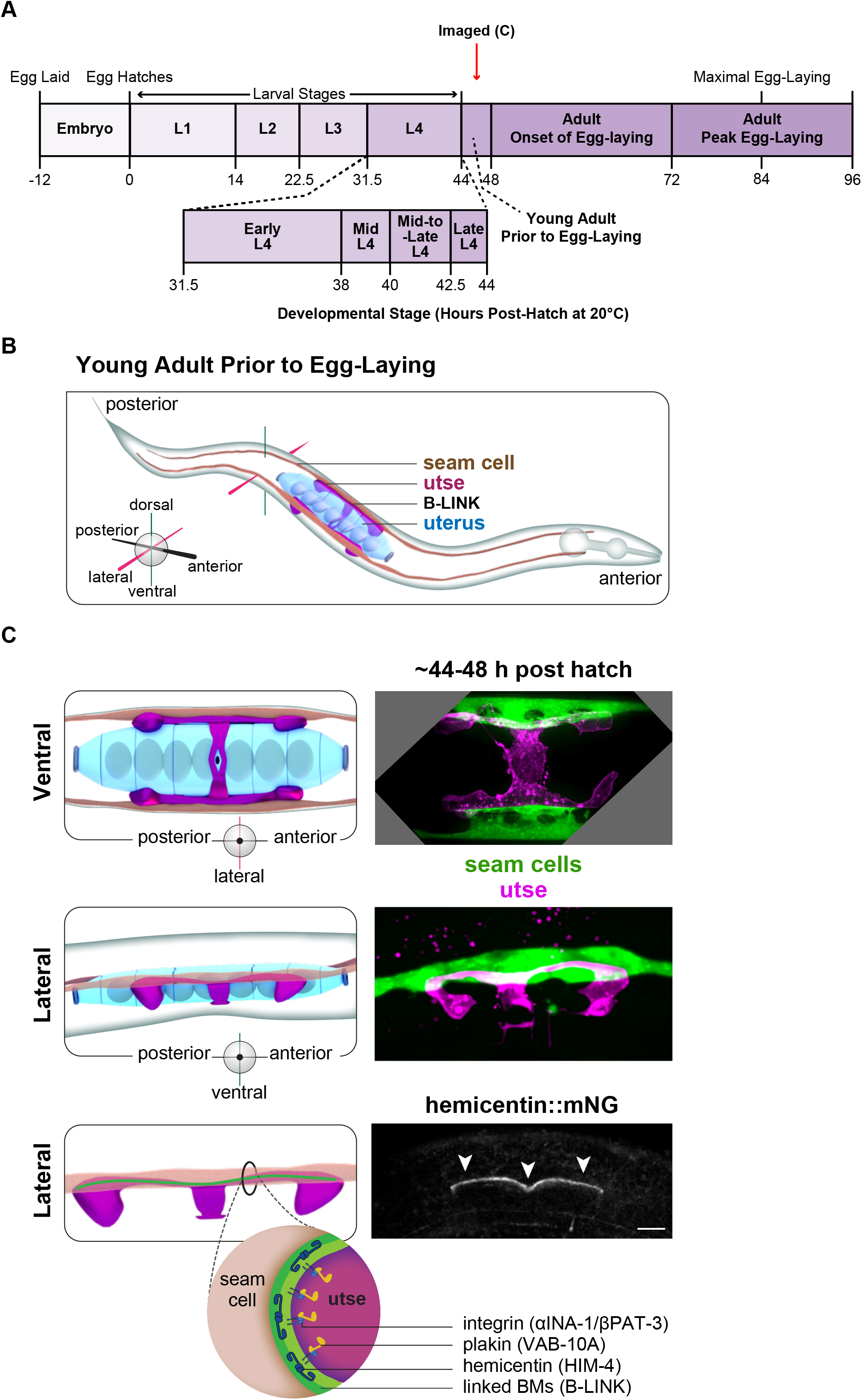
A basement membrane linkage (B-LINK) joins the uterine utse cell and epidermal seam cells in *C. elegans*. **(A)** A timeline of *C. elegans* development in hours post-hatch showing developmental stages through peak egg-laying in adulthood. **(B)** A schematic of the utse-seam B-LINK in the young adult, just prior to egg-laying. At this time, a B-LINK connects the utse (magenta) to the epidermal seam cells (brown), which run along the lateral sides of the animal (individual seam cells not shown). The utse underlies the uterus (blue). **(C)** (top) The utse (visualized with *cdh-3p::mCh::PH*, magenta) and seam cells (*scmp::GFP::CAAX*, green) from a ventral perspective and (middle) a lateral perspective. (bottom) The known B- LINK matrix component hemicentin (*C. elegans* HIM-4) visualized from a lateral perspective sits between the utse basement membrane (BM) and seam cell BM. Integrin (αINA-1/βPAT-3) and plakin (VAB-10A) are also components of the B-LINK (schematic) and function in the utse cell. Scale bar 10 µm.

**Figure 2.**
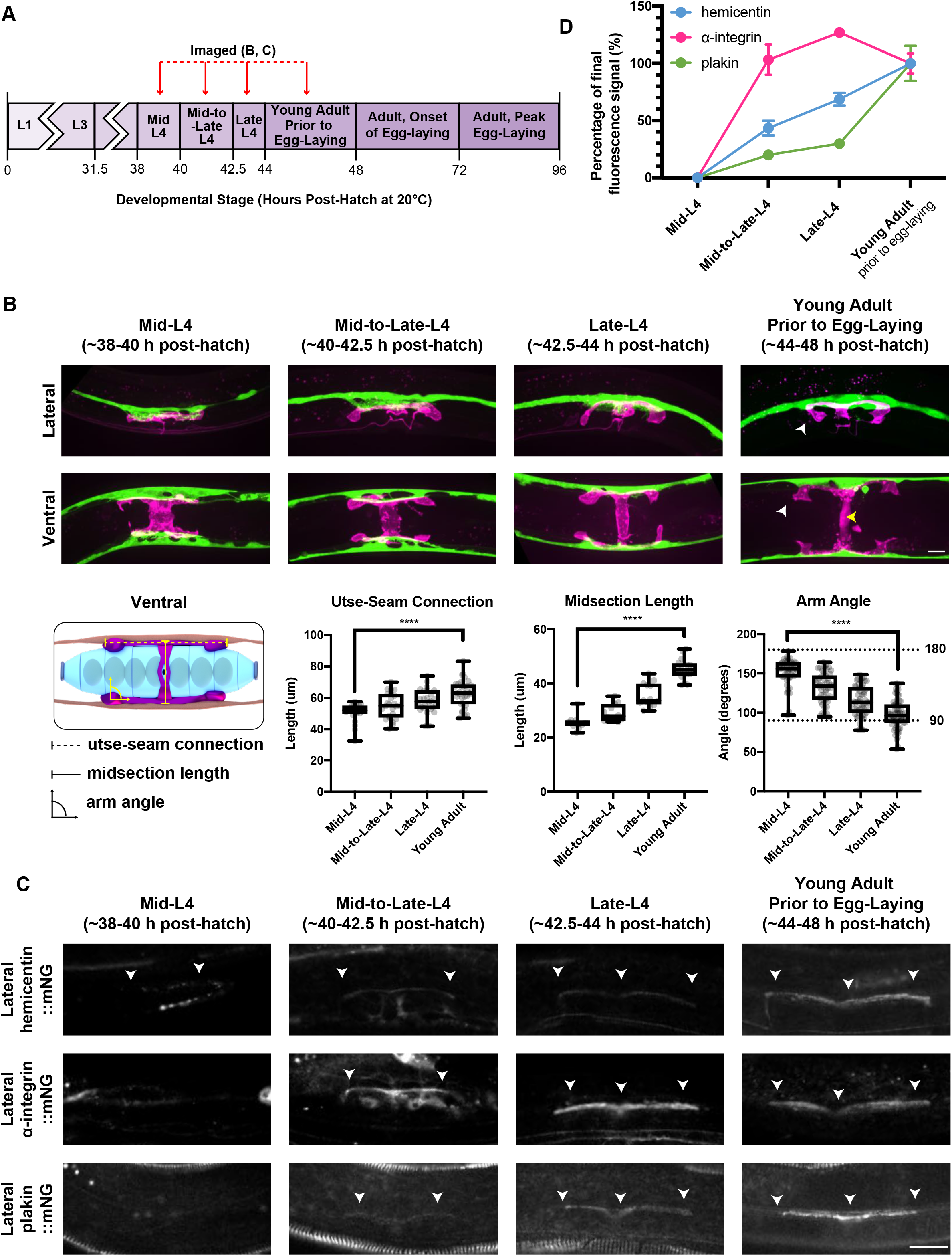
Utse and seam cell association and B-LINK formation occur during the L4 larval stage. **(A)** A schematic diagram detailing the timing of imaging. **(B)** (top) Merged fluorescence images of the utse (magenta) and seam cells (green) viewed from a lateral and ventral perspective from the mid-L4 to the young adult stage. By the young adult stage, the end of each utse arm has turned inwards away from the seam (white arrowheads) and the utse midsection is fully extended (yellow arrowhead). (bottom left) A schematic showing where measurements were taken. (bottom right) Mean length of utse-seam association (n = 10 each), mean utse midsection length (n = 10 each), and mean angle between the turned in utse arm and the seam (n = 40 each). ****P < 0.0001, one-way ANOVA followed by post hoc Tukey’s test. Box edges in boxplots depict the 25^th^ and 75^th^ percentiles, the line in the box indicates the median value, and whiskers mark the minimum and maximum values. **(C)** Localization of the known B-LINK components (white arrowheads) viewed laterally from the mid-L4 to the young adult stage. **(D)** Quantification of mean florescence intensity of each component at each timepoint as a percentage of the final mean fluorescence intensity of that component at the young adult stage. Error bars represent standard error of the mean (n = 10 all stages). Scale bars 10 µm.

To determine when the B-LINK forms to connect the utse and seam cells, we examined the localization of B-LINK components from the mid-L4 to the early adult stage using homozygous mNeonGreen (mNG) endogenous knock-in strains of hemicentin::mNG (HIM-4), α- integrin::mNG (INA-1), and plakin::mNG (VAB-10A) (Gally et al., 2016; Keeley et al., 2020) (Fig. 2 C). B-LINK components were not consistently detected at the B-LINK at the mid-L4 stage (hemicentin 5/10; integrin and plakin 0/10), but during the mid-to-late-L4 stage assembled to form a nascent B-LINK. While the α-integrin INA-1 levels plateaued during the late-L4 stage, hemicentin and plakin continued to increase until the young adult stage (Fig. 2 D). Together, these results indicate that the B-LINK begins to form during the mid-L4 stage and then extends in length and is further enriched in B-LINK components by the young adult stage.

### Perlecan, fibulin-1, type IV collagen, and papilinS enrich at the utse-seam B-LINK

In addition to hemicentin, fibulin-1 (FBL-1) also localizes to the utse-seam B-LINK, but it is unknown if fibulin-1 or any other additional matrix components have functions at this linkage (Morrissey et al., 2014; Muriel et al., 2005, 2012; Feitosa et al., 2012). To determine if other matrix components are concentrated at the utse-seam B-LINK, we examined nine endogenously tagged BM matrix components—the core BM components *γ*-laminin::mNG (LAM- 2), *α*1-type IV collagen::mNG or ::mRuby2 (EMB-9), perlecan::mNG (UNC-52), nidogen::mNG (NID-1), and type XVIII collagen::mNG (CLE-1), as well as the matricellular components papilinS::mNG (MIG-6S), papilinL::mNG (MIG-6L), spondin::mNG (SPON-1), agrin::mNG (AGR- 1), and mNG::fibulin-1 (FBL-1) (Jayadev et al., 2019; Keeley et al., 2020). *C. elegans* synthesizes two laminin heterotrimers comprising one of two *α*-chains, one *β*-chain, and one *γ*- chain; and a single triple helical type IV collagen molecule comprising two *α*1 chains and one *α*2 chain (Kramer, 2005). For simplicity, we refer to the levels and presence of *γ*-laminin::mNG (LAM-2) as laminin and *α*1-type IV collagen::mNG or ::mRuby2 (EMB-9) as type IV collagen. We performed a visual screen at the young adult stage to determine if these matrix proteins are enriched at the utse-seam B-LINK in comparison to the neighboring unlinked gonadal BM (Fig. 3). As the B-LINK is a connection between two BMs, we defined enrichment as an over two-fold higher fluorescence intensity compared to the single gonadal BM (where the utse resides). We found that perlecan, fibulin-1, and type IV collagen were significantly enriched over the gonadal BM (fold enrichment of 13.1, 8.7, and 4.7 respectively). PapilinS was also slightly enriched, although to a much lesser degree (∼2.5 fold, Fig. 3). Together, this suggests that additional matrix components enrich at the utse-seam B-LINK and could have roles in BM-BM linkage.

**Figure 3.**
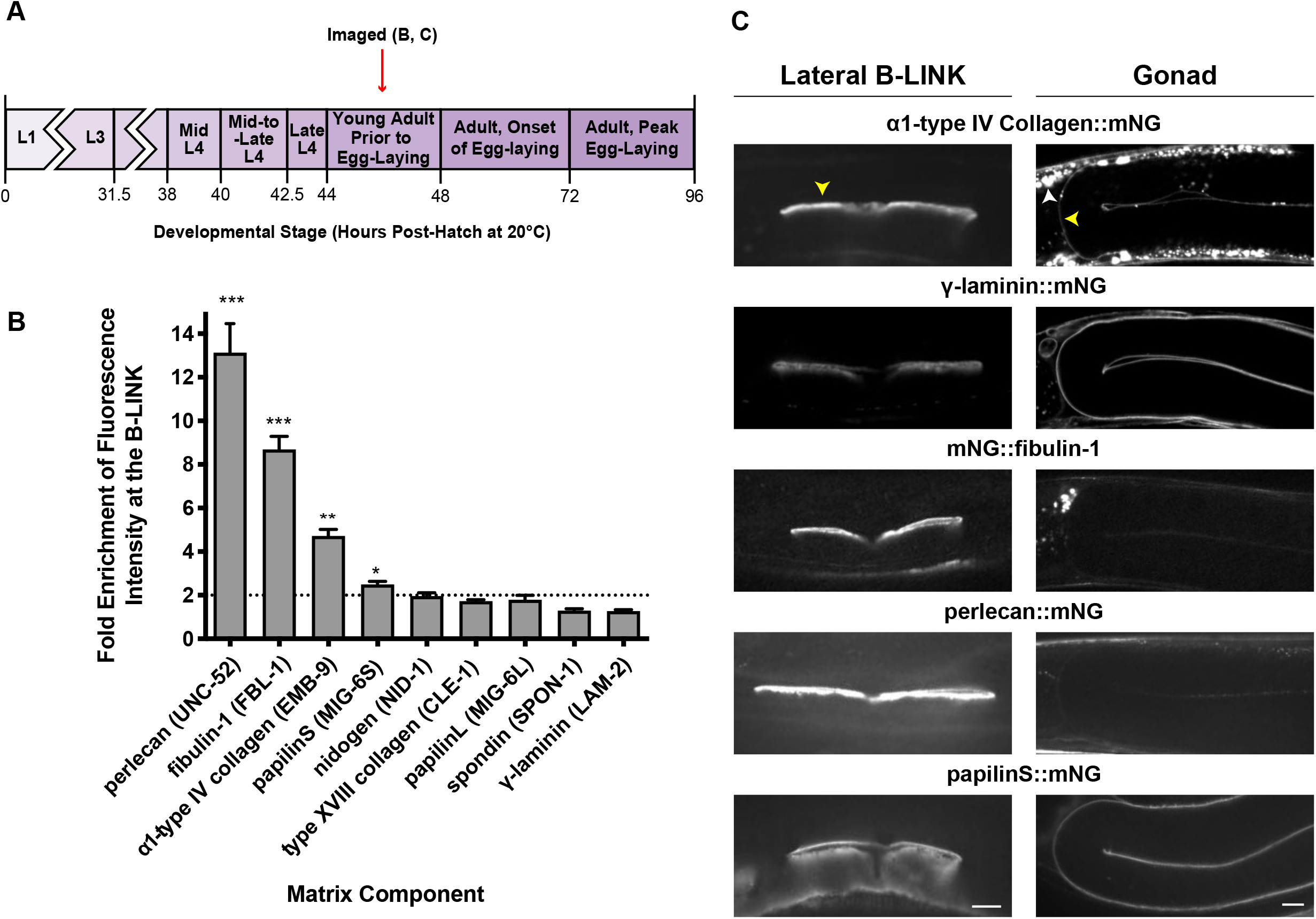
Perlecan, fibulin-1, type IV collagen, and papilinS are enriched at the B-LINK. **(A)** A schematic diagram showing when animals were imaged. **(B)** The quantification of fold enrichment of matrix component fluorescence intensity at the B-LINK compared to the gonadal BM at the young adult stage (n = 10 each). Agrin is not shown as it was undetectable at the B- LINK and in the gonadal BM (n = 10 each). ***P < 0.0001, **P < 0.001, *P < 0.01, unpaired two- tailed Student’s t-test comparing B-LINK signal to two times the gonad signal (to account for double basement membrane at the B-LINK). Error bars represent standard error of the mean. **(C)** Fluorescence images of enriched matrix components and a representative non-enriched component (*γ*-laminin::mNG) at the young adult stage at the B-LINK and in the gonadal BM. Yellow arrowheads indicate location of fluorescence intensity measurements. White arrowhead indicates *α*1-type IV collagen localized in the muscle endoplasmic reticulum where it is predominantly made. Scale bars 10 µm.

### B-LINK matrix components resist mechanical forces during egg-laying

The egg-laying neural circuit that controls egg-laying is established during the L4 stage, and vulval muscle calcium transients suggest some muscle activity may occur and generate forces on the B-LINK at this time (Ravi et al., 2018). However, it is not until the young adult stage (∼48 h post-hatch) that neuronal signals trigger regular intervals of egg-laying muscle contractions followed by expulsion of eggs from the uterus (Schafer, 2006). Loss of hemicentin or other B- LINK components can result in failure of the utse-seam B-LINK after egg-laying begins, causing the rupture (Rup) phenotype where the worm’s internal organs are extruded through the vulva (Morrissey et al., 2014; Vogel and Hedgecock, 2001). However, whether egg-laying and muscle contractions are directly responsible for the Rup phenotype has not been determined. To test this, we used a strain with optogenetically inducible muscle contraction, achieved through expression of the ultraviolet light receptor LITE-1 in body wall and egg-laying muscles (Edwards et al., 2008). Expression of LITE-1 is thought to increase calcium levels and trigger muscle contraction (Edwards et al., 2008). We placed this strain on hemicentin RNAi at the L1 stage (immediately post-hatch) and exposed worms to blue light after 60 hours (soon after the start of egg-laying) to induce powerful, extended muscle contraction and subsequent egg-laying (Fig. 4, Movie S2 and S3, RNAi knockdown efficiencies Table S2). We found that light-induced muscle contraction caused the Rup phenotype in 76% of RNAi treated animals (n = 68/90) compared to 3% of controls (n = 3/90) (Fig. 4). These observations indicate that the utse-seam B-LINK helps the reproductive tissues resist the high mechanical stress of muscle contractions and egg- laying.

**Figure 4.**
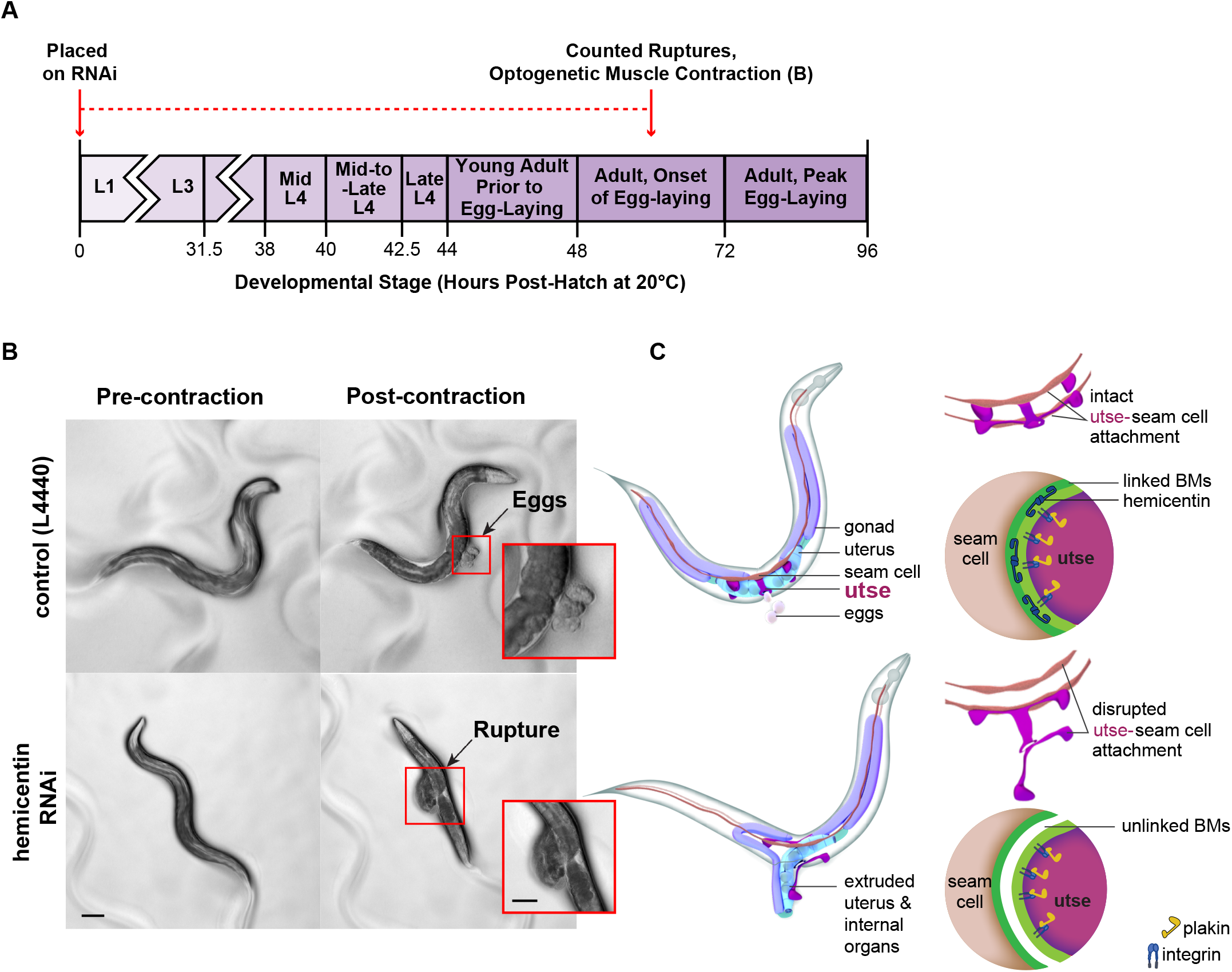
The utse-seam B-LINK requires hemicentin to withstand the mechanical forces of egg-laying. **(A)** A schematic diagram outlining the RNAi feeding and timing of optogenetic muscle contraction and imaging. **(B)** Transgenic worms expressing the ultraviolet light receptor LITE-1 in body wall and egg-laying muscles (*lite-1(ce314); ceIs37* (*myo-3p::lite-1 + myo- 3p::GFP*)) were plated on control (T444T empty vector) and hemicentin (*him-4*) RNAi and after 60 hours were exposed to 488 nm light for 7 seconds to induce muscle contractions that forced egg-laying. Images show worms both pre- and post-contraction. Regions of the animals in the red box are magnified in the insert to show egg-lay and rupture (control rupture: 3%, n = 3/90; hemicentin RNAi knockdown rupture: 76%, n = 68/90). Images are stills from supplemental video S2 (control RNAi) and S3 (hemicentin RNAi). **(C)** A schematic model of utse-seam B-LINK rupture upon loss of hemicentin. Main Image scale bar 100 µm, insert scale bar 50 µm.

We next wanted to determine if other BM components, particularly those enriched at the B- LINK, are functionally required for BM-BM linkage. To test this, we used mutants and RNAi-mediated knockdown to assess if BM component loss resulted in the Rup phenotype (Table S1). RNAi feeding was started at the L1 stage (immediately post-hatch) and animals were observed each day for the Rup phenotype until 120 hours post-hatch (past egg-laying peak). We found that loss of type IV collagen, papilinS, and fibuilin-1, which were identified in the visual screen, as well as laminin, resulted in the Rup phenotype. Notably, the frequency of rupture upon loss of fibulin-1 was low, likely as loss of fibulin-1 sharply reduces fertility and thus egg-laying (Muriel et al., 2005; Hesselson et al., 2004). Additionally, depletion of perlecan, which was also identified in the visual screen, did not cause rupture. This is likely due to the RNAi causing full muscle paralysis, as perlecan functions in anchoring muscle attachments (Bix and Iozzo, 2008; Rogalski et al., 1995; Mullen et al., 1999).

As type IV collagen, papilinS, and laminin are present in most *C. elegans* BMs, their loss could cause general BM defects that could lead to the Rup phenotype (Gordon et al., 2019; Kawano et al., 2009; Keeley et al., 2020). To test whether the Rup phenotype for these components was a result of their role as BM components or a B-LINK specific defect, we initiated RNAi knockdown at the L3 stage, ∼10 hours before the development of the B-LINK (∼30 h post-hatch). We reasoned that this late larval RNAi should reduce the later assembly of matrix components at the B-LINK but minimize depletion of components in the already assembled gonadal and seam cell BMs. RNAi against hemicentin, which specifically localizes to the B-LINK and is present at low or undetectable levels in the gonadal and seam cell BMs, was used as a positive control. We found that L3-initiated loss of hemicentin and type IV collagen produced a robust Rup phenotype (Table S1). Importantly, L3 initiated RNAi significantly decreased hemicentin and type IV collagen levels at the B-LINK (∼100% and 85%, respectively, Table S3), but type IV collagen gonadal BM levels were only slightly reduced (∼30%, Table S3), strongly suggesting the Rup phenotype for these components is due to a B-LINK role. In contrast, L3 RNAi mediated loss of laminin and papilin did not produce a Rup phenotype, suggesting that the L1 RNAi Rup phenotype resulted from defects in BM and not a B-LINK role. Consistent with this, loss of laminin and papilinS cause dramatic perturbations in gonadal BM assembly (Keeley et al., 2020; Jayadev et al., 2019).

To further test if type IV collagen and fibulin-1 are functionally required for the B-LINK, we completed optogenetic experiments on animals treated with type IV collagen or fibulin-1 RNAi and found that light-induced muscle contraction and egg-laying led to the Rup phenotype in both conditions (control RNAi, 2% rupture, n = 2/90; type IV collagen RNAi, 92% rupture, n = 85/92; fibulin-1 RNAi, 26% rupture, n = 23/90). We conclude that hemicentin, type IV collagen, and fibulin-1 (and possibly perlecan) are components of the utse-seam B-LINK and are functionally required to resist the physical stress of egg-laying.

### Hemicentin and fibulin-1 reach maximal levels prior to type IV collagen and perlecan

To investigate the roles of matrix components in B-LINK formation and function, we first examined their appearance and levels throughout B-LINK development. We found that B-LINK matrix components are undetectable or only beginning to be deposited at the mid-L4 stage (fibulin-1 and perlecan, 10/10; hemicentin, 5/10; type IV collagen, 0/10; Fig. 5 A and B). Component levels continued to increase through the mid-L4 stage and there was a robust build- up by the young adult stage. Interestingly, while hemicentin and fibulin-1 reached their maximal levels at egg-laying onset, type IV collagen and perlecan continued to enrich into the time of peak egg-laying (Fig. 5 A and B). We also assessed these component levels at the anchor cell B-LINK and found hemicentin was the only matrix component specifically enriched beneath the anchor cell (Fig. S1), indicating the utse-seam B-LINK has a more complex matrix composition. As these strains are endogenously tagged with the same mNG fluorophore, we next compared matrix component levels at the utse-seam B-LINK. In adult animals (∼72-96 h post hatch, peak egg-laying) type IV collagen and perlecan levels were ∼10-fold higher than hemicentin and ∼3- fold higher than fibulin-1 (Fig. 5 C). These findings suggest hemicentin and fibulin-1 might play an earlier role in B-LINK formation, while type IV collagen and perlecan may play a later role in stabilizing the B-LINK during adulthood when there is higher mechanical strain from egg-laying.

**Figure 5.**
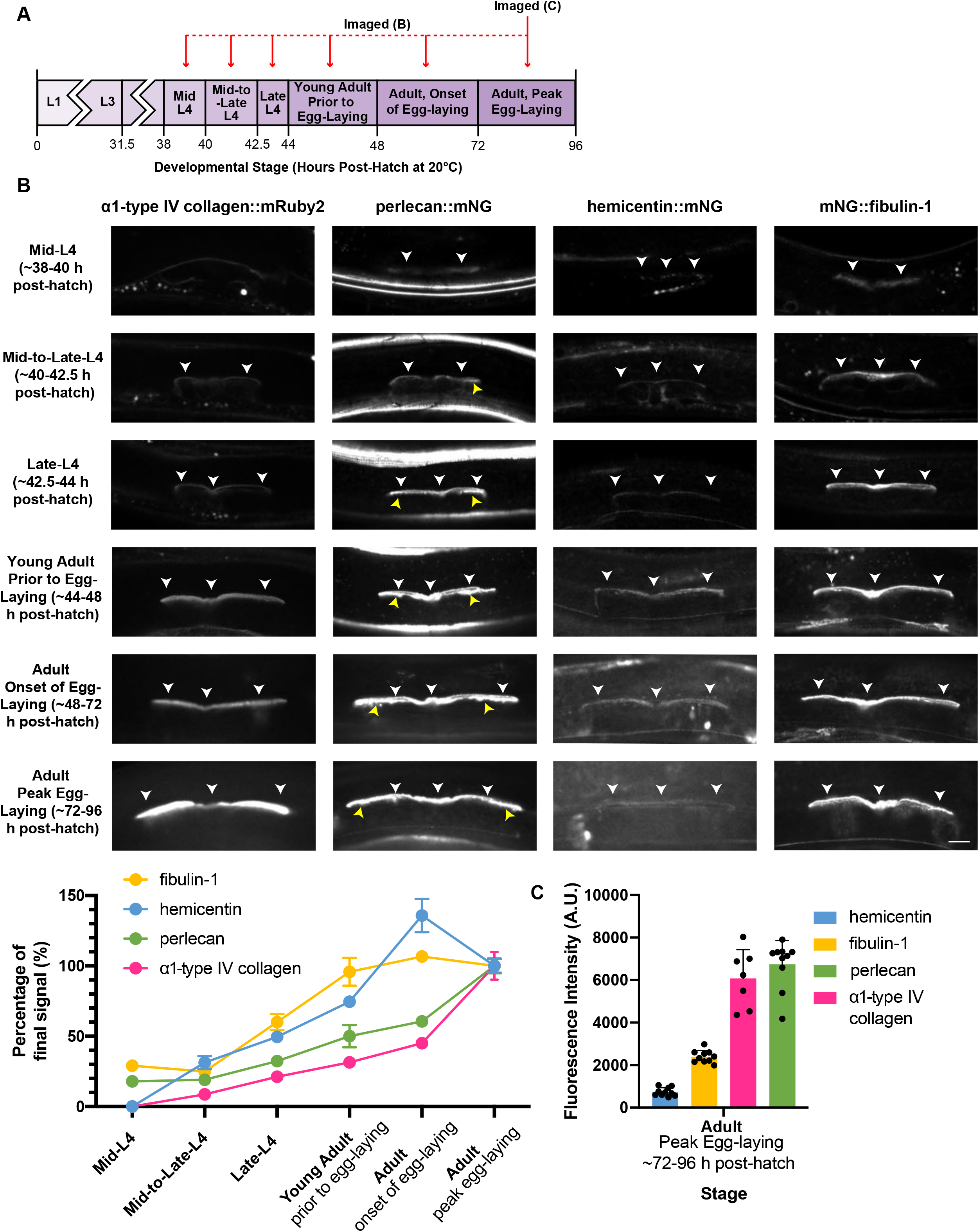
B-LINK levels of hemicentin and fibulin-1 peak earlier than type IV collagen and perlecan. **(A)** A schematic diagram delineating the timing of imaging of utse-seam B-LINK matrix components shown below. **(B)** (top) Fluorescence images of endogenous α1-type IV collagen::mRuby2, perlecan::mNG, hemicentin::mNG and mNG::fibulin-1, at the B-LINK (white arrowheads) from mid-L4 to peak egg-laying in the adult (n = 10 each). Yellow arrowheads indicate perlecan signal in the vulval muscle attachment sites below the ends of the B-LINK. (bottom) Quantification of mean florescence intensity of each B-LINK matrix component at each timepoint as a percentage of the final mean fluorescence intensity of that component in the peak egg-laying adult. Error bars represent standard error of the mean (n = 10 all stages). **(C)** Quantification of fluorescence intensity of all endogenously mNG tagged utse-seam B-LINK matrix components (imaged at the same exposure to compare levels). Error bars represent standard error of the mean. Scale bar 10 µm. A.U., arbitrary units.

### Type IV collagen and perlecan are highly stable at the B-LINK, while hemicentin and fibulin-1 are more dynamic

To gain further insight into the roles of different matrix proteins in BM-BM linkage, we assessed the stability of type IV collagen, perlecan, fibulin-1, and hemicentin at the B-LINK by performing fluorescence recovery after photobleaching (FRAP) at the young adult stage. After photobleaching, fibulin-1 and hemicentin recovered ∼50% or more of their original fluorescence intensity after 30 minutes. In contrast, type IV collagen and perlecan exhibited negligible recovery (Fig. 6 A-D). We explored type IV collagen dynamics further and found that type IV collagen recovered only ∼40% of its original fluorescence six hours after photobleaching (Fig. 6 E). This suggests that type IV collagen and perlecan are scaffolding components that stably link the utse-seam BMs together throughout adulthood, while the more dynamic hemicentin and fibulin-1 may play distinct dynamic structural roles and possibly regulatory functions at the B- LINK.

**Figure 6.**
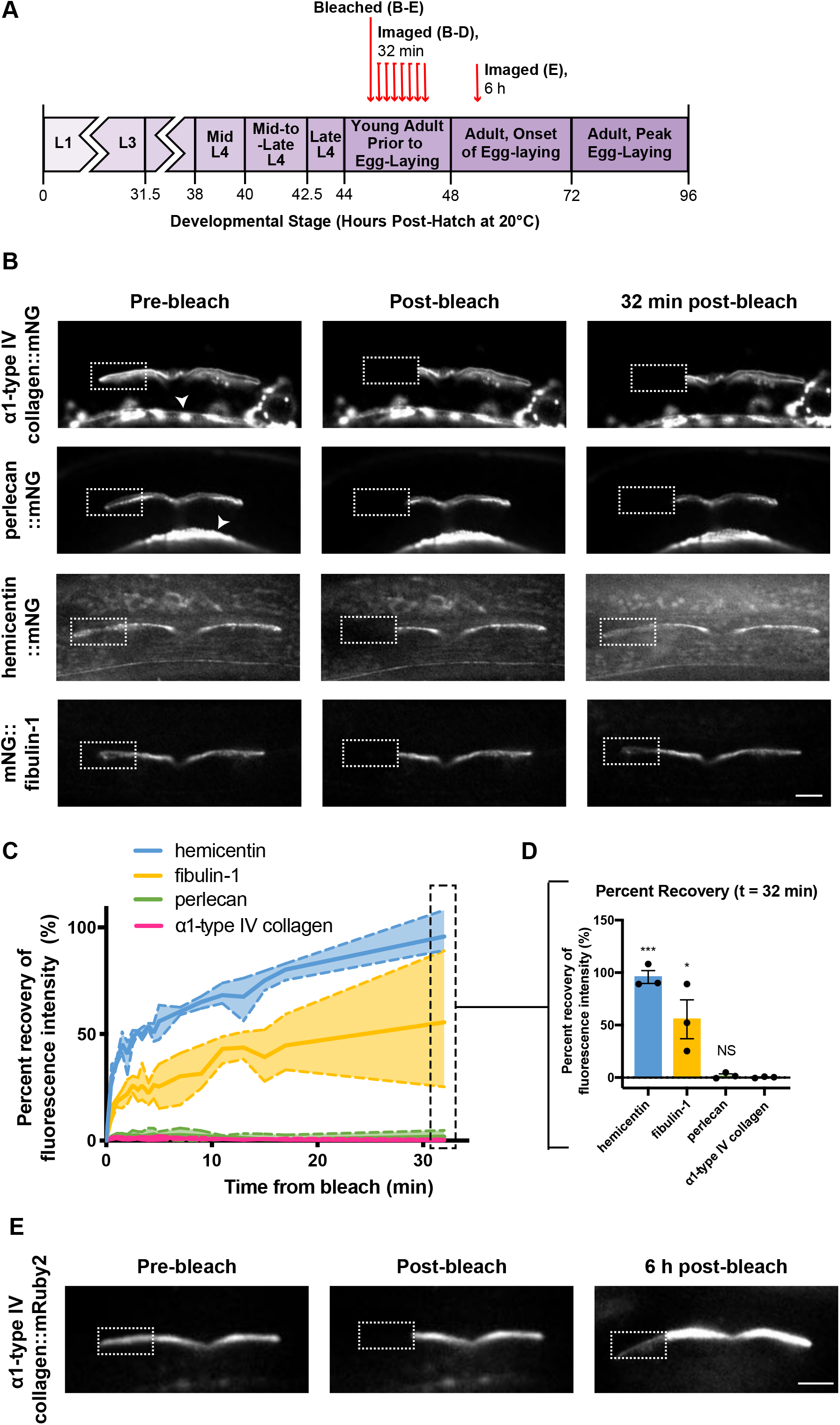
Type IV collagen and perlecan are highly stable at the utse-seam B-LINK, hemicentin and fibulin-1 are dynamic. **(A)** A schematic image showing the timing of photobleaching and recovery analysis. **(B)** Fluorescence images of *α*1-type IV collagen::mNG, perlecan::mNG, hemicentin::mNG, and mNG::fibulin-1 before photobleaching, immediately after photobleaching, and 32 minutes post photobleaching in young adult animals. Dotted white line indicates bleached region. White arrowheads indicate *α*1-type IV collagen::mNG and perlecan::mNG in the body wall muscle. **(C)** A line graph showing the fluorescence recovery of each component in the young adult B-LINK over 32 minutes (n = 3 each). Dashed lines represent the minimum and maximum values for each matrix component. **(D)** Bar graph shows the mean percent recovery of each component after 32 minutes. All components were compared with *α*1-type IV collagen. *** P < 0.001, * P < 0.01, one-way ANOVA followed by post hoc Dunnet’s test, NS non-significant. **(E)** Fluorescence images of *α*1-type IV collagen::mNG, before photobleaching, immediately after photobleaching, and six hours post photobleaching in adult animals (46 +/- 10% recovery, n = 3). Dotted white line indicates bleached region. Scale bars 10 µm.

### The utse contributes hemicentin to the utse-seam B-LINK

Most *C. elegans* BM components are secreted from body wall muscle into the extracellular fluid and then assembled on different tissues. (Clay and Sherwood, 2015). Interestingly, hemicentin is expressed by the utse (Morrissey et al., 2014; Vogel and Hedgecock, 2001), suggesting that hemicentin, and possibly other B-LINK matrix components, could be contributed locally. To test this, we performed L1 RNAi targeting hemicentin, type IV collagen, fibulin-1, and perlecan using a strain where only the uterine tissue is sensitized to RNAi and examined if this resulted in the Rup phenotype. We found that uterine-specific RNAi against only hemicentin caused rupture, indicating that the utse contributes hemicentin to support the utse-seam B-LINK (Table S4). We also tested whether the seam cell contributes B-LINK components, but seam cell-specific RNAi did not cause any ruptures (Table S4). We conclude that hemicentin is provided by the utse, while type IV collagen, fibulin-1 and perlecan appear to be recruited to the B-LINK from the extracellular fluid.

### Hemicentin is required for assembly of other B-LINK matrix components

The local production of hemicentin at the B-LINK suggested it might have an organizational role. Consistent with this possibility, hemicentin has previously been reported to promote fibulin-1 assembly (Muriel et al., 2005). To confirm hemicentin’s role in fibulin-1 assembly and determine whether assembly of type IV collagen and perlecan might also be dependent on hemicentin, we examined their levels at the utse-seam B-LINK at the young adult stage (48 h post-hatch, just before egg-laying) after loss of hemicentin (L1-mediated RNAi, 100% knockdown, Table S2). Consistent with a crucial organizational role at the B-LINK, hemicentin was required for the robust assembly of all B-LINK matrix components (Fig. 7). In addition, loss of every component caused a reciprocal increase in hemicentin, suggesting feedback between hemicentin and other B-LINK components (Fig. S2). Fibulin-1 loss caused minimal changes in other B-LINK components, while type IV collagen loss caused a modest reduction of fibulin-1, but a significant reduction in perlecan (Fig. S3). These experiments suggest that hemicentin plays a crucial role in the assembly of other B-LINK matrix components, and that there is feedback between hemicentin and other B-LINK components.

**Figure 7.**
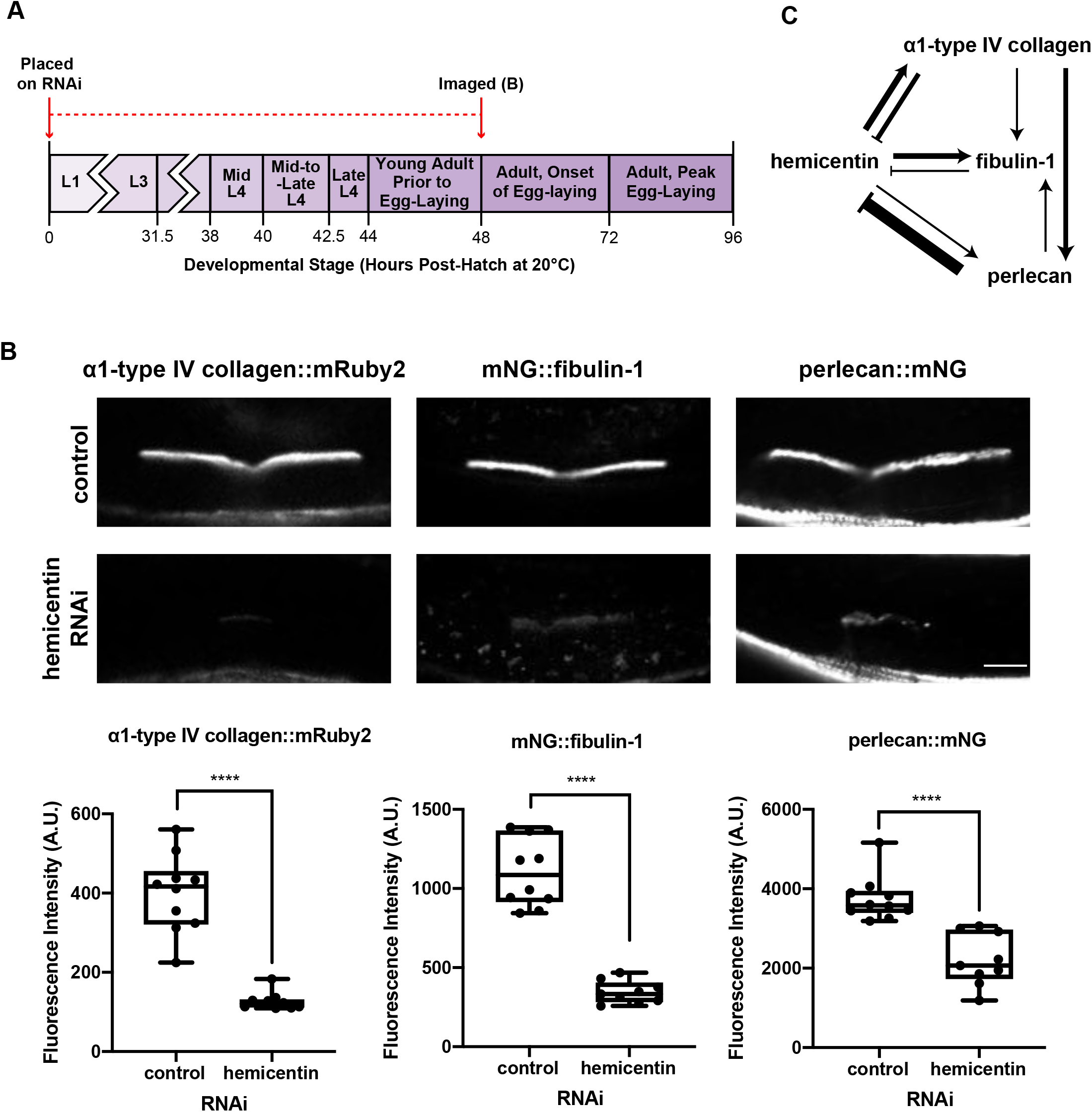
Utse-seam B-LINK matrix component assembly is dependent on hemicentin. **(A)** A schematic diagram outlining timing of hemicentin RNAi and imaging of utse-seam B-LINK matrix components. **(B)** (top) Fluorescence images of *α*1-type IV collagen::mRuby2, mNG::fibulin-1, and perlecan::mNG on control (T444T empty vector, n = 10 each) and hemicentin (*him-4*, n = 10 each) RNAi. (below) Mean fluorescence intensity of each component quantified. ****P < 0.0001, one-way ANOVA followed by post hoc Dunnet’s test. Box edges in boxplots depict the 25^th^ and 75^th^ percentiles, the line in the box indicates the median value, and whiskers mark the minimum and maximum values. Scale bar 10 µm. A.U., arbitrary units. **(C)** A model of component recruitment to the utse-seam B-LINK. See Fig. S2 and S3 for further details.

### Hemicentin and fibulin-1 facilitate initial B-LINK formation and type IV collagen is required for maintenance during egg-laying

Peak levels of hemicentin and fibulin-1 occur earlier at the B-LINK than the later accumulation of type IV collagen and perlecan. To test whether this might indicate temporal roles in B-LINK formation and maintenance, we knocked down each B-LINK component using L1 RNAi and imaged the utse and seam cells at the mid-L4, late-L4, and adult onset of egg-laying stages (Table S2). Defects in the B-LINK were indicated by a lack of contact between the utse and seam cells (utse-seam gaps). Fibulin-1 loss caused the most penetrant early defect at the mid- L4 stage, indicating fibulin-1 has the earliest role at the utse-seam B-LINK (Fig. 8 A-C). Importantly, contact between the utse and seam at the early-L4 stage was unimpaired after fibulin-1 loss (10/10 animals), suggesting fibulin-1 loss does not alter the initial utse and seam cell contact. We found loss of hemicentin also caused an early B-LINK defect, with the utse occasionally pulling off from the seam at the mid-L4 stage, but more significantly by the late-L4 (Fig. 8 B and C). Interestingly, fibulin-1 deposition was consistently present at the mid-L4 stage (Fig. 5 B; 10/10 animals), while hemicentin was inconsistently present (Fig. 5B; 5/10 animals). Knockdown of hemicentin did not severely affect early fibulin-1 deposition (only ∼20% reduction in fibulin-1 levels, Fig. 8 D), although fibulin-1 became completely reliant on hemicentin by the late-L4 (Fig. 8 D). These data indicate that fibulin-1 has an early hemicentin-independent role in mediating utse-seam B-LINK formation, but by the late L4 becomes dependent on hemicentin. Loss of type IV collagen did not affect the utse-seam B-LINK through the L4 stage (Fig. 8 B and C), and only caused utse-seam separation in the adult at the initiation of egg-laying (Fig. 8 B and C). These results strongly suggest that type IV collagen strengthens the B-LINK so that it can withstand higher mechanical forces during egg-laying. Perlecan RNAi did not cause utse-seam separation at any timepoint (Fig. 8 B and C), likely because its loss led to complete paralysis and the absence of mechanical stress on the B-LINK.

**Figure 8.**
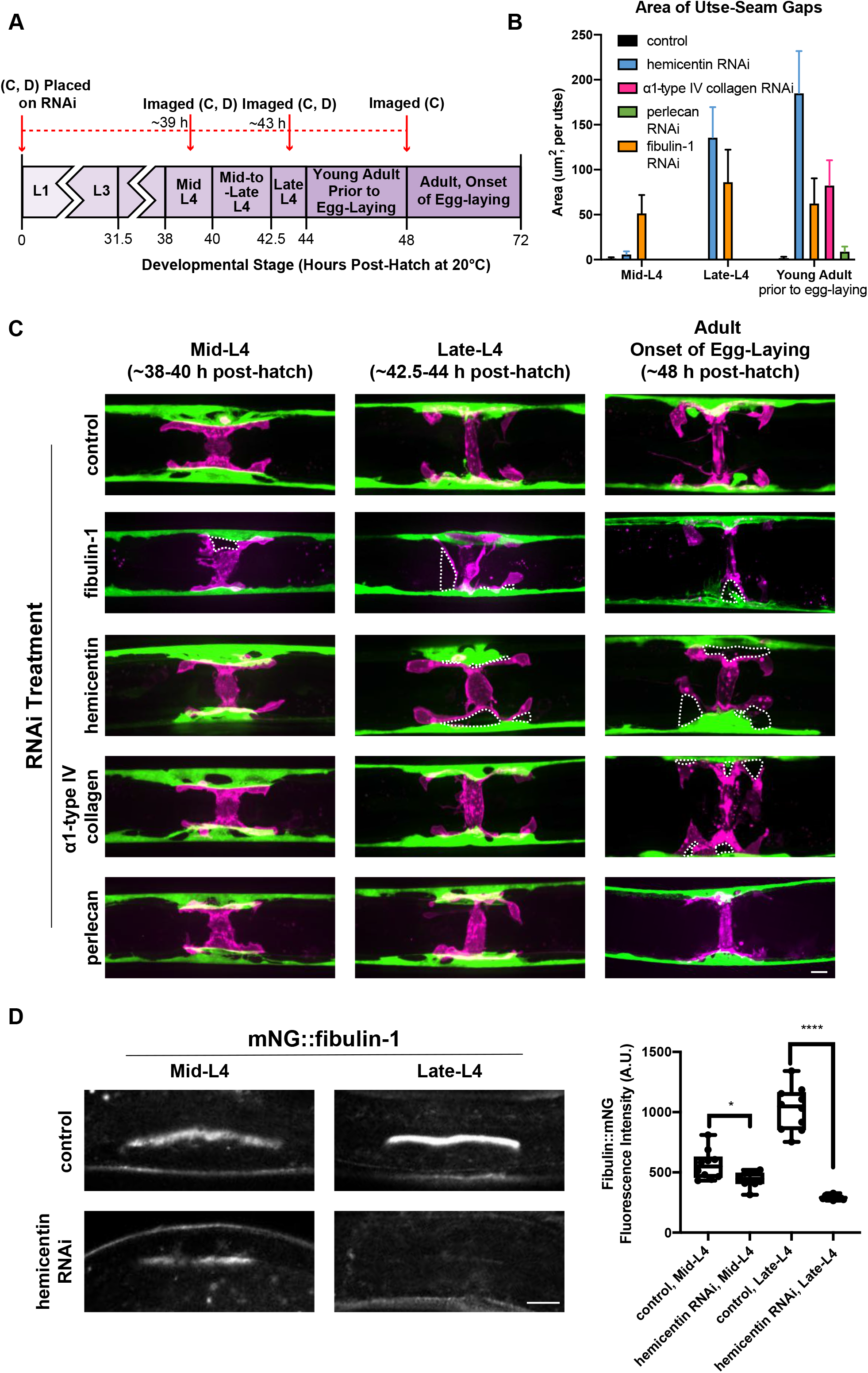
Hemicentin and fibulin-1 initiate utse-seam BM-BM linkage, while type IV collagen maintains the B-LINK during egg-laying. **(A)** A schematic diagram outlining timing of RNAi initiation and imaging of utse-seam cell association. (B) Quantification of the total utse- seam cell gap area per utse cell imaged in (C). Error bars represent standard error of the mean. (C) Ventral fluorescence images of the utse (magenta) and seam cells (green) at stages shown on control (L4440/T444T empty vectors, n = 10 each) and after RNAi-mediated reduction of B- LINK matrix components (n = 10 each). Regions where the utse has pulled away from the seam (utse-seam gaps) indicating defects in the B-LINK are highlighted with white dotted lines. (D) (left) Fluorescence images of mNG::fibulin-1 on control (T444T empty vector, n = 10 each) and hemicentin (*him-*4, n = 10 each) RNAi at the mid-L4 and late-L4 stages. (right) Quantification of mean fluorescence intensity at each stage on both control and hemicentin RNAi. Box edges in boxplots depict the 25^th^ and 75^th^ percentiles, the line in the box indicates the median value, and whiskers mark the minimum and maximum values. ****P < 0.0001, *P < 0.01 unpaired two-tailed Student’s t-test. Scale bars 10 µm. A.U., arbitrary units.

### Hemicentin and fibulin-1 have early roles in B-LINK formation, but not maintenance

We next examined the role of hemicentin, fibulin-1 and type IV collagen during B-LINK maintenance. We first assessed hemicentin by initiating RNAi-knockdown at the early-L4 stage (∼5 h before B-LINK formation), the mid-L4 stage (the start of B-LINK formation), the mid-to-late-L4 stage (∼5 h after B-LINK starts to form), or the young adult stage (∼10 h after B-LINK starts to form) and scoring for the Rup phenotype (Fig. 9 A, Table S5). We found that knockdown of hemicentin at the early-L4 stage or mid-L4 stage (the start of B-LINK formation) caused rupture, while knockdown of hemicentin at the young adult stage did not cause rupture despite a complete loss of detectable hemicentin (Fig. 9 B; Table S5). This suggests hemicentin is required during the early stages of utse-seam B-LINK formation, but not for B-LINK maintenance. To confirm this, we placed animals on hemicentin RNAi at the young adult stage (48 h post-hatch, ∼10 h after B-LINK initiation) and imaged the utse and seam cells 24 hours later (72 h post-hatch, just before peak egg-laying adult stage). This later RNAi plating did not cause separation of the utse and seam cells, indicating there was no B-LINK defect (Fig. 9 C). These results provide compelling evidence that hemicentin has an early role during utse-seam B-LINK formation, but is dispensable after the B-LINK is established.

**Figure 9.**
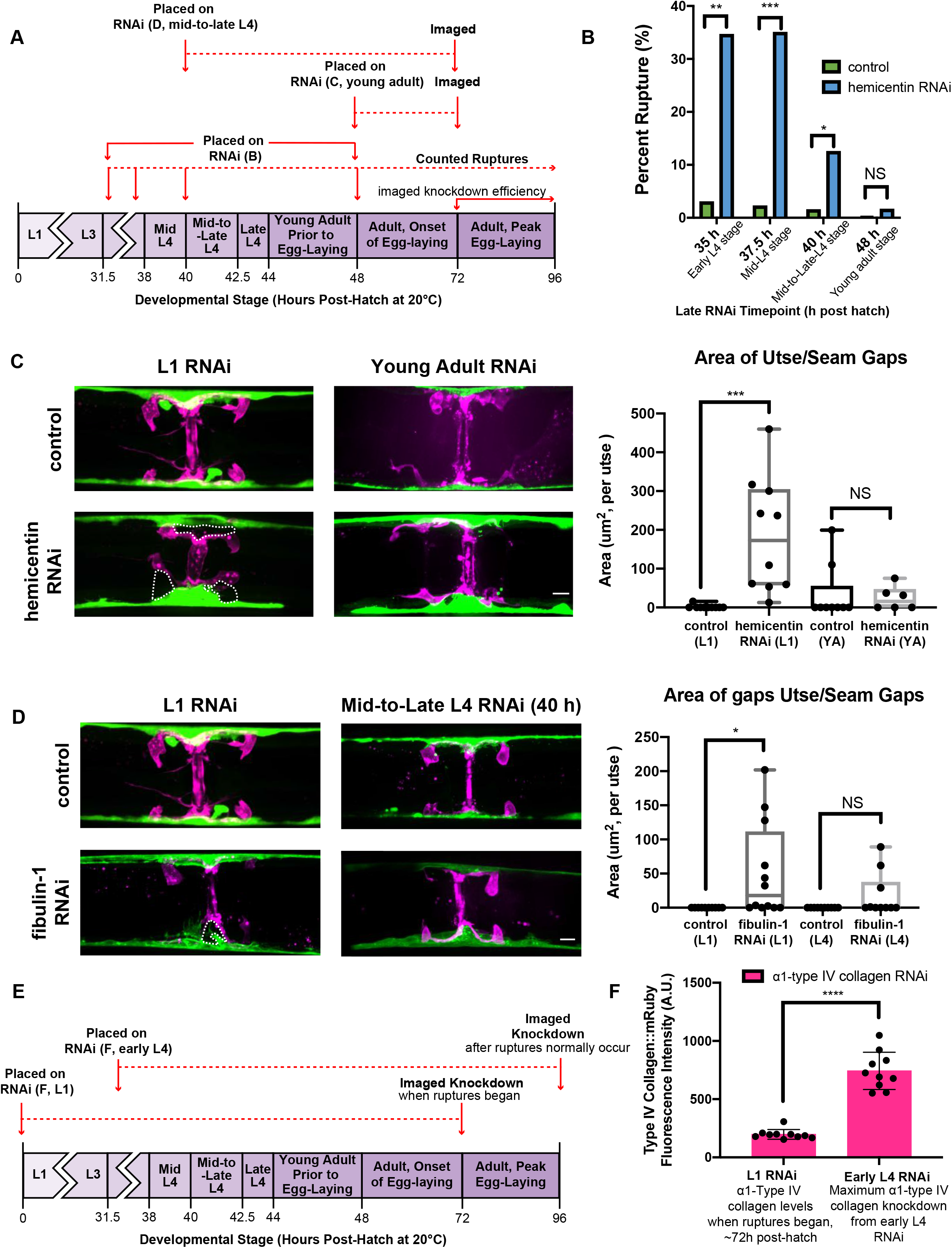
Hemicentin is not required for later maintenance of the utse-seam B-LINK. **(A)** A schematic diagram detailing the timing of RNAi initiation and utse-seam connection analysis in panels (B-D). **(B)** The percent of animals ruptured after 120 h on control (T444T empty vector) or hemicentin RNAi that was started at either the early L4 stage (control n = 33, hemicentin n = 26), the mid-L4 stage (control n = 25, hemicentin n = 40), the late-L4 stage (control n = 26, hemicentin n = 48), or the young adult stage (control n = 243, hemicentin n = 127). ***P < 0.001, **P < 0.01, *P < 0.05, Fisher’s Exact Tests, NS, not significant. **(C)** Ventral fluorescence images of the utse (magenta) and seam (green) on hemicentin RNAi started at either the L1 stage or at the young adult (YA) stage (L1 data shown previously in Fig. 8 B and C). Regions where the utse has pulled away from the seam (utse-seam gaps) indicating defects in the B-LINK are highlighted with white dotted lines and quantified on the right (n = 10 each). Box edges in boxplots depict the 25^th^ and 75^th^ percentiles, the line in the box indicates the median value, and whiskers mark the minimum and maximum values. ***P < 0.001, unpaired two-tailed Student’s t- tests, NS, not significant. **(D)** Ventral fluorescence images of the utse and seam on fibulin-1 RNAi started at the L1 stage or at the mid-to-late-L4 stage (L1 data shown previously in Fig. 8 B and C). Utse-seam gaps (dotted lines) indicating defects in the B-LINK are quantified on the right (n = 10 each). Box edges in boxplots depict the 25^th^ and 75^th^ percentiles, the line in the box indicates the median value, and whiskers mark the minimum and maximum values. *P < 0.05, unpaired two-tailed Student’s t-tests, NS, non-significant. **(E)** A schematic image detailing timing of RNAi knockdown of *α*1-type IV collagen and analysis of its levels in panel (F). **(F)** Quantification of *α*1-type IV collagen::mRuby2 levels at the B-LINK when Rup phenotypes began after L1-initiated *α*1-type IV collagen RNAi compared to a1-type IV collagen::mRuby2 levels at the B-LINK after 120 h on *α*1-type IV collagen RNAi started at the early L4 stage. Error bars represent standard error of the mean. ****P < 0.0001, unpaired two-tailed Student’s t-test. Scale bars 10 µm. A.U., arbitrary units.

We next investigated fibulin-1. As fibulin-1 is required for fertility, which affects the penetrance of the Rup phenotype, we directly examined the utse and seam cells. To determine if fibulin-1 was required after its early role in B-LINK formation, we placed animals on fibulin-1 RNAi at the young adult stage (∼10 h after B-LINK initiation) to assess utse-seam separation 24 hours later. RNAi platings at this stage, however, did not result in sufficient fibulin-1 protein knockdown (Table S6). Thus, RNAi was used to knock down fibulin-1 at the mid-to-late-L4 stage and the utse and seam cells were examined 32 hours later at the onset of egg-laying. Although RNAi knockdown of fibulin-1 was greater than 50% (Table S6), fibulin-1 loss did not cause utse-seam separation (Fig. 9 D). In contrast, early loss of fibulin-1 (L1 RNAi, also ∼50% knockdown, Table S2) resulted in separation of the utse and seam cells, indicating a B-LINK defect (Fig. 9 D). These results strongly suggest that like hemicentin, fibulin-1 has a specific early role in B-LINK formation, but not in maintenance during egg-laying.

As type IV collagen appeared to be required for maintenance of the mature B-LINK, we hypothesized that RNAi knockdown of type IV collagen beginning at the late-L4 and young adult stages would cause rupture during peak egg-laying. However, no significant ruptures were observed (late L4, 1/35; young adult, 1/246). We hypothesized that since type IV collagen is highly stable at the B-LINK, the lack of late ruptures could be because type IV collagen loss was not significant enough to disrupt the B-LINK. To test this, we knocked down type IV collagen using L1 RNAi and measured type IV collagen levels at the B-LINK at rupture onset (∼72 h post hatch), thus determining the threshold collagen levels when the B-LINK begins to fail. We then compared this to type IV collagen levels after later RNAi platings (Fig. 9 E and F). Although L4 RNAi platings caused ∼75% knockdown of type IV collagen (at least 60 h on RNAi, Fig 9 E), the remaining type IV collagen levels were > 2-fold higher than type IV collagen levels when ruptures were first observed (Fig. 9 F). To allow more time for knockdown, we plated worms on type IV collagen RNAi at the young adult stage (48 h post-hatch) and scored for rupture through the adult decline in egg-laying stage (end of day 4 adult) and found that ruptures were significantly higher than controls at this later timepoint when minimal egg-laying is occurring (control 2%, 2/90; type IV collagen RNAi 16%, 14/90; fisher’s exact test P < 0.01). Together, this indicates that hemicentin and fibulin-1 initiate utse-seam BM-BM attachment, while type IV collagen is highly stable and functions to maintain the utse-seam B-LINK during the mechanically active time of egg-laying.

### Hemicentin and fibulin-1 levels dramatically decline at the B-LINK during egg-laying

As our experiments indicated early roles for fibulin-1 and hemicentin in mediating utse-seam B- LINK formation and later roles for type IV collagen and perlecan in B-LINK maintenance, we examined the localization of these components during B-LINK maintenance. We quantified component levels starting at the adult peak egg-laying stage (72 h post-hatch) as well as two other later timepoints when egg-laying is gradually declining because of germline aging (120 h post-hatch/∼day 3 adult and 168 h post-hatch/∼day 4 adult) (Fig. 10 A) (Kocsisova et al., 2019). Both hemicentin and fibulin-1 levels significantly decreased throughout adulthood and reached low and, in the case of hemicentin, nearly undetectable levels (Fig. 10 B and C). In contrast, perlecan and type IV collagen levels, were either maintained (perlecan, 10 animals per stage) or continued to increase (type IV collagen, 10 animals per stage), consistent with a specific role in B-LINK maintenance.

**Figure 10.**
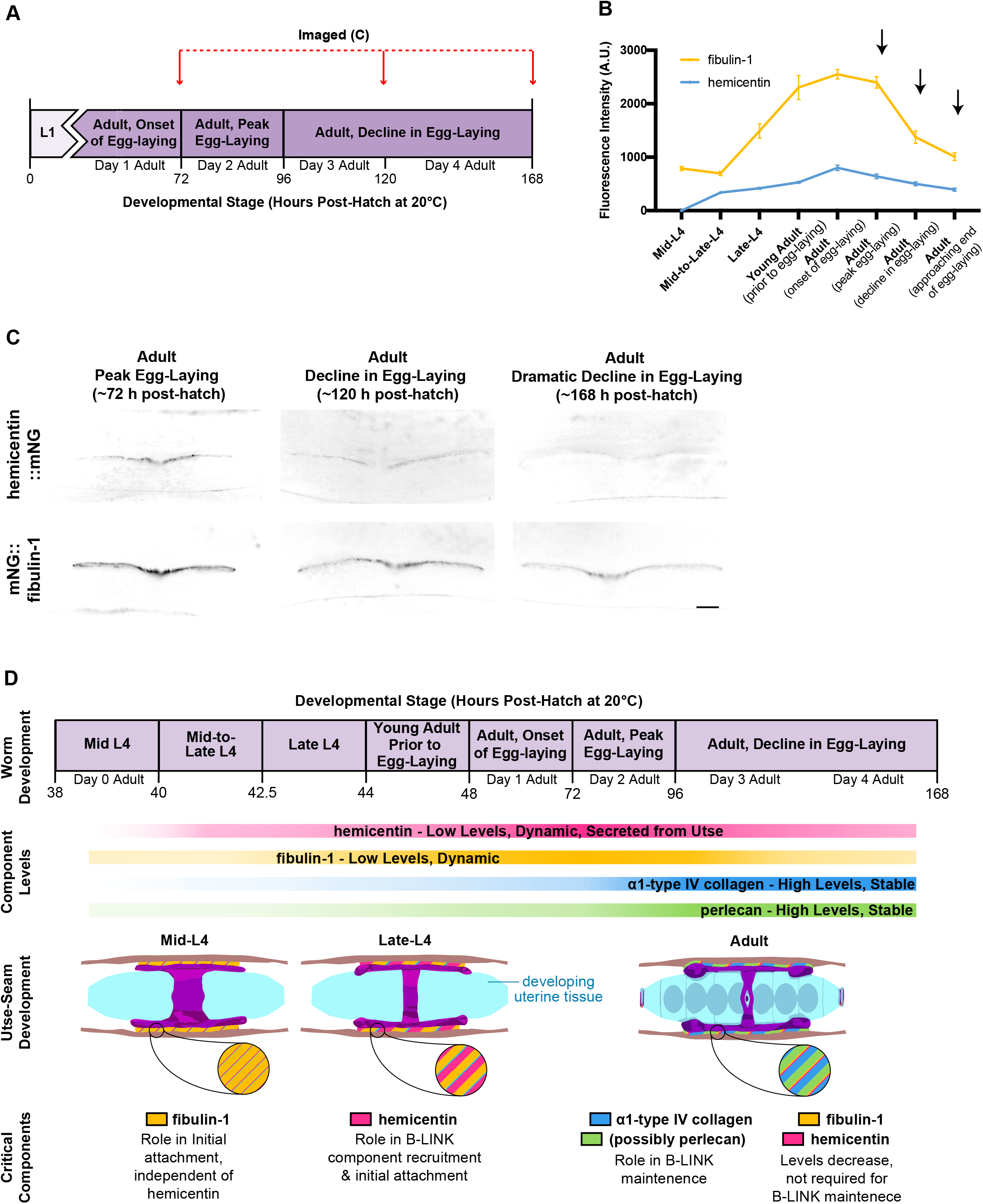
Hemicentin and fibulin-1 decline at the utse-seam B-LINK during egg-laying. **(A)** A schematic diagram outlining when animals were imaged for presence of mNG::fibulin-1 and hemicentin::mNG at the utse-seam B-LINK. **(B)** Quantification of mean florescence intensity of fibulin-1 and hemicentin from mid-L4 through adult decline in egg-laying stage (mid-L4 through onset of egg-laying data shown previously in Fig. 5 B). Black arrows indicate fluorescence time points shown in panel **(C)**. Error bars represent standard error of the mean (n = 10 all stages). (C) Fluorescence images of hemicentin::mNG and mNG::fibulin-1 at the B-LINK at the adult peak egg-laying, adult decline in egg-laying 120 h/∼day 3 adult, and adult decline in egg-laying 168 h/∼day 4 adult stages. Colors are inverted to improve visibility. **(D)** A model for assembly and role of different matrix components at the utse-seam B-LINK. Scale bars 10 µm. A.U., arbitrary units.

## Discussion

Basement membranes (BMs) enwrap and separate most tissues, allowing them to slide along each other (Brown, 2011). However, at specific locations, including the lung alveoli, kidney glomeruli, and astrocytes in the blood-brain barrier, adjacent tissues link their BMs to stabilize tissues and build specialized organs (Keeley and Sherwood, 2019). Due to the challenge of dissecting complex tissue interactions, however, the mechanisms that mediate BM-BM tissue attachment are poorly understood. We previously characterized a transient BM-BM linkage (B- LINK) that facilitates anchor cell invasion through linked uterine and vulval BMs and is composed of the matricellular matrix protein hemicentin, the matrix receptor integrin, and the cytolinker plakin (Morrissey et al., 2014). We also discovered that hemicentin, integrin, and plakin are components of a stable B-LINK that connects the multi-nucleated uterine utse cell to seam epidermal cells for the lifetime of the animal. This utse-seam BM-BM linkage has been proposed to maintain the uterus within the animal during egg-laying (Vogel and Hedgecock, 2001). It was unknown, however, when the utse-seam B-LINK forms, if it contains additional matrix components to mediate a more robust BM-BM linkage, and whether it directly functions to resist the high mechanical stress of egg-laying.

Using live imaging and endogenous mNeonGreen tagged hemicentin, integrin, and plakin, we discovered that the nascent B-LINK begins to form at the mid-L4 stage and enriches in B-LINK components by the young adult stage, just prior to egg-laying. Using LITE-1 optogenetic stimulation of body wall and egg-laying muscle contractions (Edwards et al., 2008) on hemicentin depleted animals (weakened utse-seam B-LINK), we also found that egg-laying causes uterine tissue prolapse. These experiments establish that the utse-seam B-LINK forms prior to egg-laying and that it functions to resist mechanical forces associated with muscle contraction and egg-laying.

Through screening and RNAi depletion, we discovered the utse-seam B-LINK has additional matrix components not found at the anchor cell B-LINK. These include the matricellular protein fibulin-1, and core BM proteins type IV collagen and perlecan. Similar to hemicentin, loss of fibulin-1 and type IV collagen led to uterine prolapse during egg-laying. Because of its function in anchoring muscle attachments, however, perlecan loss eliminated egg-laying muscle contractions (Rogalski et al., 1995), thus impeding the determination of its functional role at the B-LINK. Similar to hemicentin, each of these components appeared at the nascent B-LINK between the mid-L4 and the late-L4 stages and enriched into the young adult stage. We also discovered that hemicentin is contributed to the B-LINK locally from the utse and plays a key role in promoting the assembly of all other B-LINK matrix components as the B-LINK matures. Notably, previous transmission electron microscopy studies have observed electron dense extracellular matrix material localized between the utse and seam cell BMs that is lost in hemicentin null mutants (Vogel and Hedgecock, 2001). These observations are consistent with this material being composed of hemicentin and other B-LINK matrix components whose assembly hemicentin promotes. Interestingly, we also found that loss of fibulin-1, type IV collagen and perlecan caused a reciprocal increase in hemicentin, suggesting feedback mechanisms set the precise levels of matrix proteins at the B-LINK. Close association of hemicentin with fibulins and collagens have been observed in both *C. elegans* and vertebrates (Feitosa et al., 2012; Lin et al., 2020; Muriel et al., 2005). The physical binding partners of hemicentins, however, are poorly understood (Zhang et al., 2021). Studies in *C. elegans* have shown hemicentin promotes the assembly of fibuilin-1 at several extracellular sites and requires EGF repeats 4 and 5 of fibulin-1, suggestive of a direct interaction (Muriel et al., 2005, 2012). How hemicentin promotes type IV collagen assembly, however, is unclear. We found that perlecan assembly during B-LINK maturation is largely dependent on collagen and prior work has shown that perlecan binds type IV collagen (Hohenester and Yurchenco, 2013), suggesting hemicentin promotes perlecan enrichment indirectly through collagen.

Using timed RNAi depletion, photobleaching, and quantitative fluorescence analysis, we also determined distinct properties and temporal requirements for matrix components at the utse-seam B-LINK (summarized in Fig 10 D). We found that hemicentin and fibulin-1 have dynamic associations, with residence times of less than thirty minutes, and that both proteins reached peak levels prior to egg-laying. In contrast, perlecan and type IV collagen reached peak levels later during egg-laying. Consistent with their early deposition, hemicentin and fibulin-1 loss caused B-LINK disruption and utse-seam splitting at the mid and late L4 stages before type IV collagen levels ramped up. Although the B-LINK does not resist the mechanical forces of egg-laying at this time, vulval and possibly uterine muscle contractions appear to occur in the L4 stage (Ravi et al., 2018), which would place some mechanical stress on the B-LINK. Notably, once type IV collagen began to build up during egg-laying, hemicentin and fibulin-1 were no longer required to maintain the B-LINK, and their endogenous levels dramatically decreased. This supports the idea of a specific early role for fibulin-1 and hemicentin in mediating the initial stages of BM-BM connection. It is likely that hemicentin and fibulin-1 have independent roles in linking the BMs, as loss of fibulin-1 led to defects prior to loss of hemicentin. Further, we’ve previously shown that hemicentin alone promotes BM-BM linkage under the anchor cell (Morrissey et al., 2014), and hemicentin-1 in zebrafish mediates fin fold BM-BM linkage (Carney et al., 2010). Hemicentin-2 and fibulin-1 also function together in zebrafish to facilitate a transient somite and epithelial BM-BM connection (Feitosa et al., 2012). These observations suggest that hemicentin and fibulin-1 function as a dynamic matrix that can rapidly assemble to stabilize BM-BM linkages, withstand modest mechanical forces, and help promote the assembly of a more stable BM-BM tethering matrix.

In contrast to hemicentin and fibulin-1, we found that type IV collagen and perlecan are both highly stable at the utse-seam B-LINK and enrich to maximal levels after the onset of egg-laying. Consistent with this later build-up, type IV collagen depletion only began to cause utse-seam B-LINK defects at the onset of egg-laying, suggesting it functions to bear high mechanical loads. Supporting this idea, the triple helix of collagen provides matrices with high tensile strength (Fidler et al., 2018). The role of collagen as a stabilizing component of BM-BM linkages may be conserved, as in mice and humans the collagen α3α4α5(IV) trimer localizes between the podocyte and glomerular BMs—fused BMs that resist strong hydrostatic pressure during blood filtration (Suleiman et al., 2013). Further, mutations in α3α4α5(IV) lead to Alport syndrome, a kidney disease that can result in kidney failure (Naylor et al., 2020). In patients with Alport syndrome, the glomerular BM often splits into distinct podocyte and endothelial BMs and it has been unclear why loss of α3α4α5(IV) collagen leads to BM separation (Naylor et al., 2020; Rumpelt, 1987). Our results suggest that type IV collagen may help stably link the podocyte and endothelial BMs. Consistent with this idea, the α3α4α5(IV) trimer is thought to have greater mechanical strength then that of the ubiquitously expressed α1α1α2(IV) trimer (Naylor et al., 2020). In addition, the collagen α3α4α5(IV) trimer is upregulated during glomerulogenesis and replaces the α1α1α2(IV) trimer at the time of BM-BM fusion (Abrahamson, 1985; Abrahamson et al., 2009). Whether this collagen also requires an earlier BM bridging matrix is unknown. Loss of the two vertebrate hemicentins do not affect the glomerular BM in mice, however, fibulin-1 is present within the glomerular BM and might compensate for hemicentin loss (Lin et al., 2020; Naylor et al., 2020). Notably, The α3α4α5(IV) trimer is also present in the ear and eye, which are sites of BM-BM linkage that are also affected in Alport syndrome (Naylor et al., 2020; Keeley and Sherwood, 2019). Thus, type IV collagen may have a broad and important role in stably maintaining other BM-BM linkages.

Nearly 20 different BM-BM linkages have been observed that connect diverse tissues (Keeley and Sherwood, 2019; Gao et al., 2017; Welcker et al., 2021). Although the molecular nature of most of these connections remain elusive, the composition and structure of characterized BM-BM linkages is flexible. For example, the fibulin-1/hemicentin matrix that connects the pharyngeal and muscle BMs in *C. elegans* is 7-9 µm long, while the fibulin-1/hemicentin at the utse-seam B-LINK span a distance of less than 0.1 µm (Vogel and Hedgecock, 2001). Matrix composition can also be distinct, as the anchor cell B-LINK harbors hemicentin, but lacks the fibulin-1, perlecan, and type IV collagen found at the utse-seam B-LINK. Yet, important commonalities in BM-BM linkages are also emerging, including the importance of hemicentins and fibulin-1 in *C. elegans* B-LINKs, zebrafish fin fold and somite-epidermal BM-BM connections, and a role in stabilizing fine interdigitations of tendon at myotendinous junctions (Morrissey et al., 2014; Vogel and Hedgecock, 2001; Welcker et al., 2021; Muriel et al., 2005; Feitosa et al., 2012; Carney et al., 2010; Suleiman et al., 2013). Further commonalties include the role of type IV collagen in bridging the glomerular BM and utse-seam B-LINK BMs (Suleiman et al., 2013), where collagen may help resist high mechanical stress. Together, these studies suggest that BM-BM linkages can be tailored for the unique spatial, temporal and mechanical needs of tissues. Understanding these linkages will not only increase our knowledge of tissue structure and function, but also help inform therapies to repair BM-BM connections in Alport syndrome and other pathologies affecting these ubiquitous, yet poorly studied tissue connections.

## Acknowledgements

We thank S. G. Payne and A. Garde for comments on the manuscript and members of the Sherwood laboratory for helpful discussions. We thank A. Kawska (info@illuscientia) for her work on schematics. C.A.G. was supported by F31HL156438 and D.R.S., C.A.G., D.P.K., W.R.L., K.P., R.J., and Q.C. were supported by R35GM118049 and R21OD028766. The authors declare no competing financial interests.

## Author contributions

D.P. Keeley, C.A. Gianakas, and D.R. Sherwood conceived the project. C.A. Gianakas designed, completed, and analyzed all experiments with the following additional contributions: Q. Chi built RNAi constructs used in Figs. 4, 7, 8, 9, Fig. S2-4, Table S1-6; K. Park acquired and analyzed the utse/seam timecourse data used in Fig. 1 C and Fig. 2 A and collected the anchor cell data shown in Fig. S1; W. Ramos-Lewis acquired and analyzed the B-LINK component timecourse shown in Fig. 10 B and C; D.P. Keeley acquired a portion of the rupture data shown in Table S1; R. Jayadev built the RNAi-sensitized strains used in Fig. 8-9, Fig. S2-4, and Table S1, S2, S3, and S6. R. Jayadev also completed the uterine and hypodermal specific RNAi experiments in Table S4. C.A. Gianakas prepared the figures. C. A. Gianakas and D.R. Sherwood wrote the initial manuscript. C.A. Gianakas, D.R. Sherwood, D.P. Keeley, K. Park, W. Ramos-Lewis, and R. Jayadev reviewed and edited the manuscript.

## Methods

### Worm handling and strains

Worms were grown under standard conditions on NGM plates seeded with OP50 *Escherichia coli* at 16°C, 18°C, or 20°C (Stiernagle, 2006). Strains used in this study are listed in Table S7.

### Cloning of RNAi into T444T vector

For RNAi against *fbl-1*, and *him-4*, new RNAi clones were generated in the T444T RNAi vector (Sturm et al., 2018). Briefly, PCR fragments corresponding to existing *fbl-1* and *him-4* RNAi clones in the Vidal (Rual et al., 2004) and Ahringer (Kamath et al., 2003) RNAi libraries respectively were amplified and inserted into SacII and HindIII-digested T444T vector by Gibson assembly. Forward and reverse primers used to amplify these fragments were: *fbl-1,* 5’- CGATGAATTCGAGCTCCACCGCGGATGCATGTGACTCTGGTACAGA-3’ and 5’ GAGGTCGACGGTATCGATAAGCTTGTTCAACGAGTCGTGAATATAG-3’; *him-4*, 5’- CGATGAATTCGAGCTCCACCGCGGAACCAACAATTTCGTGGCTCAAAG-3’ and 5’- GAGGTCGACGGTATCGATAAGCTTTTCAGCCAATTTGATAGGGCAAGT-3’. Constructs were transformed into HT115 competent cells.

### RNA Interference

All RNAi experiments were performed using the feeding method (Rual et al., 2004; Kamath et al., 2003; Timmons et al., 2001), and with the exception of *fbl-1* and *him-4*, all RNAi constructs were obtained from the Vidal and Ahringer libraries. RNAi bacterial cultures were grown in selective media (1:1000 ampicillin) for 24 h at 30°C and then for an additional hour after the addition of 1mM Isopropyl b-D-1-thiogalactopyranoside (IPTG) to induce dsRNA expression. RNAi plates were prepared by spreading a 1:1 mixture of 1M IPTG and 100mg/ml ampicillin (18μl each) on NGM agar plates. Plates were then seeded with RNAi bacterial cultures and left at room temperature overnight to allow for further induction and drying. For RNAi initiated at the L1 larval stage, synchronized L1 worms were placed on RNAi plates and allowed to feed between 24 and 120 h at 20°C depending on the experiment. The L4440 and T444T empty vectors were used as negative controls.

For RNAi initiated at later developmental stages, worms were synchronized and plated on OP50, left to grow for ∼30, 35, 40, 45, or 48 h at 20°C, then transferred to RNAi plates. Briefly, worms were washed off of OP50 plates into 1mL of M9 buffer and collected in 15 mL conical tubes, pelleted, and transferred into a non-stick polycarbonate 1.5 mL tube. Worms were washed 4x by centrifugation at 2000rpm for 60 seconds, aspirating supernatant, and resuspending in M9 buffer. After the second wash, tubes were placed on a rocking incubator for 10-15 minutes to allow worms to pass any bacteria remaining in the gut. Prior to the fourth and final wash, worms were transferred to a fresh 1.5 mL tube. After the final wash, worms were resuspended in M9 buffer and plated on RNAi-feeding plates.

To improve RNAi knockdown efficiency for the *fbl-1* RNAi, we used worms harboring a null mutation in *lin-35* [*lin-35(n745)]* (Lehner et al., 2006). Knockdown levels were quantified and are shown in Table S2, S3, S5, and S6. Box plots for all quantified knockdown experiments are in Figure S4.

Tissue specific RNAi experiments were performed as previously described using *rrf-3*(*pk1426*) II; *qyIs102 [fos-1ap::RDE-1, myo-2p::GFP]*; *qyIs10 [lam-1p::lam-1::GFP]* IV*; rde-1 (ne219)* V; *qyIs24 [cdh-3p::mCherry::PLCdPH]* worms for uterine- specific RNAi and *rde-1(ne219)* V; *kzIs9[pKK1260(lin-26p::nls::gfp)*, *pKK1253(lin-26p::rde-1)*, *pRF4(rol-6)]* worms for hypodermal-specific RNAi (Qadota et al., 2007; Tabara et al., 1999; Hagedorn et al., 2009). Briefly, these worms harbor a null mutation in *rde-1* which is required for RNAi sensitivity (Qadota et al., 2007; Tabara et al., 1999). To restore RNAi sensitivity to the tissue of interest, a functional copy of RDE-1 is specifically expressed in the tissue of interest allowing for tissue specific RNAi knockdown.

### Imaging

Confocal Images were acquired on a Zeiss AxioImager microscope equipped with a Yokogawa CSU-10 spinning disc confocal controlled by Micromanager or Metamorph software using a Zeiss 40x Plan-APOCHROMAT 1.4NA oil immersion objective or 100x Plan-APOCHROMAT 1.4NA oil immersion objective and a Hamamatsu Orca-Fusion sCMOS camera or ImageEM EMCCD camera. Worms were mounted on 5% noble agar pads containing 0.01 M sodium azide for imaging for all experiments except for Fig. 6, where worms were immobilized using 100nm polystyrene beads (Polysciences cat. #64010). For lateral images of B-LINK components we acquired z-stacks at 0.37 μm intervals spanning the entirety of the B-LINK. For lateral and ventral images of the utse and seam we acquired z-stacks spanning the top half of the utse (lateral) or the full depth of the utse (ventral).

For ventral images, slides were made by pouring agar onto a vinyl record to make agar pads with negative replicas of the record channels (Zhang et al., 2008). Worms were then added onto the record slides in M9, moved into the channels, and cover slipped (Port City Diagnostics, Inc, #M2000-10). The cover slip was then gently pushed to rotate the animals in the channels to a ventral view for imaging.

Rupture movies were acquired on a Zeiss Axio Zoom V16 stereo fluorescence microscope controlled by Zen 3.2 software using a 3x objective and an Axiocam digital camera. Movies were acquired using the Zen 3.2 movie recorder.

### Fluorescence Recovery After Photobleaching

For fluorescence recovery after photobleaching (FRAP) experiments, photobleaching was performed using an iLas^2^ FRAP system from BioVision equipped with an Omicron Lux 60mW 405nm continuous wave laser and controlled with MetaMorph software. For all FRAP experiments, worms were immobilized using 100nm polystyrene beads (Polysciences cat. #64010). Regions of interest were selected with the freehand ROI tool. One half of the uterine- epidermal junction was photobleached at 10% laser power. The number of repetitions needed to achieve complete photobleaching varied by strain and was determined experimentally. Animals were rescued after photobleaching to confirm they were able to recover and continue developing.

### Image analysis, processing, and quantification

Raw images were quantified in Fiji 2.0 (Schindelin et al., 2012). B-LINK fluorescence intensity measurements were acquired by taking the mean fluorescence intensity of a 3-pixel wide line drawn through the left side of the B-LINK in a single confocal z-slice. Background intensity values were obtained by taking a line measurement in a region with no visible fluorescence signal. Images of the utse and seam markers were acquired as z-stacks and are displayed in figures as max projections. All other images are single slices. All images of each strain within each experiment were acquired using identical settings. Fluorescence intensity values for the adult timepoints in Figure 10 B and C were normalized to data in Figure 5 B based on the overlapping adult peak egg-laying timepoint. 3D isosurfaces of the utse and B-LINK hemicentin were created from confocal z-stacks and exported using Imaris 7.4. Figures were constructed using Adobe Illustrator (CC 2021) and graphs were exported from GraphPad Prism 8.

### Scoring of ruptured through vulva (Rup) phenotype

To score for the Rup phenotype, ∼30-80 worms were plated on RNAi at the L1, L3, L4, or young adult stages as described above. After 24 h on RNAi, the number of worms on the plate was recorded. At least once every 24 h for the next 120 h, plates were visually screened for ruptured worms. Ruptured animals were subtracted from the total number of starting animals and picked off the plate so as not to double count them. After 120 h the percentage of ruptured animals was calculated and compared to controls using Fisher’s exact test.

For the long-term collagen RNAi rupture experiment, worms were synchronized and plated on RNAi at the young adult stage as previously described. Plates were then scored for ruptures as above for seven days (168 h). Each day, non-ruptured worms were picked onto fresh RNAi plates to avoid starvation and remove progeny. After seven days the percentage of ruptured animals was calculated and compared to controls using Fisher’s exact test.

For experiments using the optogenetically induced muscle contraction strain KG1271, worms were synchronized and plated on RNAi at the L1 stage as described above and fed for 72 h. Plates of animals were placed on the Axio Zoom V.16 where they were exposed to 488nm light for ∼7 seconds. After light induced muscle contraction, the number of eggs expelled by the animal was recorded and the animal was scored as positive or negative for the Rup phenotype.

### Statistical Analysis

Statistical analysis was performed in GraphPad Prism 8. Distribution of data was assessed for normality using the Shapiro-Wilk test. For comparisons of mean fluorescence intensities between two populations, we used an unpaired two-tailed Student’s *t*-test. To compare mean fluorescence intensities between three or more populations, we performed a one-way ANOVA followed by either a post hoc Dunnett’s or Tukey’s multiple comparison test. To compare percent rupture between two populations, we performed a Fisher’s Exact Test. All graphs were prepared in GraphPad Prism. Figure legends indicate sample sizes, statistical tests used, and p-values.

### Online Supplemental Material

Fig. S1, related to Fig. 5, shows the identified B-LINK matrix components that are enriched at the utse-seam B-LINK at the B-LINK underneath the anchor cell. Fig. S2, related to Fig. 7, shows fluorescence intensity of hemicentin (HIM-4::mNG) on control (L4440/T444T empty vectors) and corresponding B-LINK component RNAi (*α*1-type IV collagen, perlecan, and fibulin- 1). Fig. S3, also related to Fig. 7, shows fluorescence intensity of other B-LINK components (*α*1- type IV collagen::mRuby2, mNG::fibulin-1, and perlecan::mNG) on control (L4440/T444T empty vectors) and corresponding B-LINK component RNAi (hemicentin, *α*1-typeIV collagen, perlecan and fibulin-1). Figure S4, related to figures 4, 7, 8, and 9, shows box plots of all RNAi knockdown experiments completed with B-LINK component RNAi at various timepoints. Table S1, related to Fig. 4, shows percent rupture of all screened matrix components. Table S2, related to figures 4, 7, 8, and 9, shows percent knockdown for L1 RNAi experiments. Table S3, related to table S1, shows percent knockdown for L3 RNAi experiments. Table S4, related to table S1, shows rupture screening with the uterine specific and hypodermal specific RNAi. Table S5, related to Fig. 9, shows percent knockdown for L4 RNAi experiments where ruptures were scored. Table S6, related to Fig. 9, shows percent knockdown for L4 RNAi experiments where the utse and seam were imaged. Table S7 lists all strains used in this study.

---

Figure S1. **Utse-seam B-LINK components at the anchor cell B-LINK (A)** A schematic diagram indicating when imaging was conducted. **(B)** Florescence images (left) and corresponding differential interference contrast (DIC) image (right) of hemicentin::mNG, mNG::fibulin-1, α1-type IV Collagen::mRuby2, and perlecan::mNG underneath the anchor cell (yellow arrowhead) when a B-LINK connects the uterine and epidermal BMs. Hemicentin is enriched underneath the anchor cell (yellow arrowhead) compared to the neighboring BM (white arrowhead), while other matrix components are not enriched. White arrows indicate the anchor cell in DIC images. Scale bar 5 µm.

Figure S2. **Dynamic feedback between hemicentin and other B-LINK matrix components**. **(A)** A schematic figure showing the timing of RNAi treatment and hemicentin::mNG imaging. **(B)** (left) Lateral view fluorescence images of hemicentin::mNG on control (empty vector, n = 10), *α*1-type IV collagen, perlecan, and fibulin-1 RNAi (n = 10 each). Note increased hemicentin signal in other tissues on perlecan RNAi. (right) Mean fluorescence intensity measurements of hemicentin on each RNAi. ****P < 0.0001, ** P < 0.01, *P < 0.05, one-way ANOVA followed by post hoc Dunnet’s test, unpaired two-tailed Student’s t-test for fibulin-1 RNAi. Box edges in boxplots depict the 25^th^ and 75^th^ percentiles, the line in the box indicates the median value, and whiskers mark the minimum and maximum values. Scale bar 10 µm. A.U., arbitrary units. **(C)** A model of matrix interactions that dictate their composition at the utse-seam B-LINK. Also shown in Fig. 7 C.

Figure S3. **Dependency of B-LINK matrix components on each other at the B-LINK. (A)** A schematic diagram showing the timing of RNAi knockdown and analysis of utse-seam B-LINK matrix component levels. **(B-D)** Mean fluorescence intensity of B-LINK matrix components *α*1- type IV collagen::mRuby2, mNG::fibulin-1, and perlecan::mNG on control (L4440/T444T empty vectors) and corresponding B-LINK matrix component RNAi. The role of hemicentin in recruiting other B-LINK matrix components is shown in Fig. 7 and Fig. S2. *lin-35(n745)* was used for fibulin-1 RNAi to sensitize worms for more efficient knockdown. Box edges in boxplots depict the 25^th^ and 75^th^ percentiles, the line in the box indicates the median value, and whiskers mark the minimum and maximum values. ****P < 0.0001, *P < 0.05, one-way ANOVA followed by post hoc Dunnet’s test, unpaired two-tailed t-test for fibulin-1 RNAi. A.U., arbitrary units.

Figure S4. **RNAi Knockdown efficiencies of utse-seam B-LINK matrix components. (A-D)** Quantification of knockdown efficiency for B-LINK matrix component RNAi started at the L1, L3, and L4 stage. ****P < 0.0001, unpaired two-tailed t-tests, NS, non-significant. Box edges in boxplots depict the 25^th^ and 75^th^ percentiles, the line in the box indicates the median value, and whiskers mark the minimum and maximum values. A.U., arbitrary units.

Video S1. **Hemicentin located at the utse-seam B-LINK.** Movie shows a 3-Dimensional reconstruction of hemicentin (visualized with hemicentin::GFP) localized at the ends of the utse cell (visualized with cdh-3p::mCh::moeABD) that contact the seam cells (not shown). Note, the utse marker cdh-3p::mCh::moeABD is expressed in four groups of vulval muscles that extend below the cross-bar of the utse. Related fluorescence images in Figure 1. Scale bar labelled in video.

Video S2. **Optogenetically induced muscle contraction causes egg-laying.** A transgenic worm expressing the ultraviolet light receptor LITE-1 in body wall and egg-laying muscles (*lite- 1(ce314); ceIs37* (*myo-3p::lite-1 + myo-3p::GFP*)) grown 60 hours on control T444T empty vector RNAi and then exposed to 488 nm light for 7 seconds to induce extended body wall and egg-laying muscle contractions and forced egg-laying. Stills from movie are shown in Fig 4 B. Video was collected and is displayed at 31.5 frames/second. Scale bar labelled in video.

Video S3**. Optogenetically induced muscle contraction causes uterine rupture after loss of hemicentin.** A transgenic worm expressing the ultraviolet light receptor LITE-1 in body wall and egg-laying muscles (*lite-1(ce314); ceIs37* (*myo-3p::lite-1 + myo-3p::GFP*)) grown 60 hours on hemicentin RNAi and then exposed to 488 nm light for 7 seconds to induce body wall and egg-laying muscle contractions. The animal shown undergoes uterine rupture during forced egg-laying. Stills from movie are shown in Fig. 4 B. Video was collected and is displayed at 31.5 frames/second. Scale bar labelled in video.

Table S1. **Rupture screening of basement membrane matrix components**

Table S2. **L1 RNAi knockdown efficiencies**

Table S3. **L3 RNAi knockdown efficiencies**

Table S4. **Rupture screening of basement membrane matrix components**

Table S5. **L4 RNAi knockdown efficiencies, rupture experiments**

Table S6. **L4 RNAi knockdown efficiencies, utse-seam imaging**

Table S7. **Strain information**

## Notes

### Competing Interest Statement

The authors have declared no competing interest.

