## Supplemental Figures & Tables for "Hemicentin mediated type IV collagen assembly strengthens juxtaposed basement membrane linkage"

### Figure S1

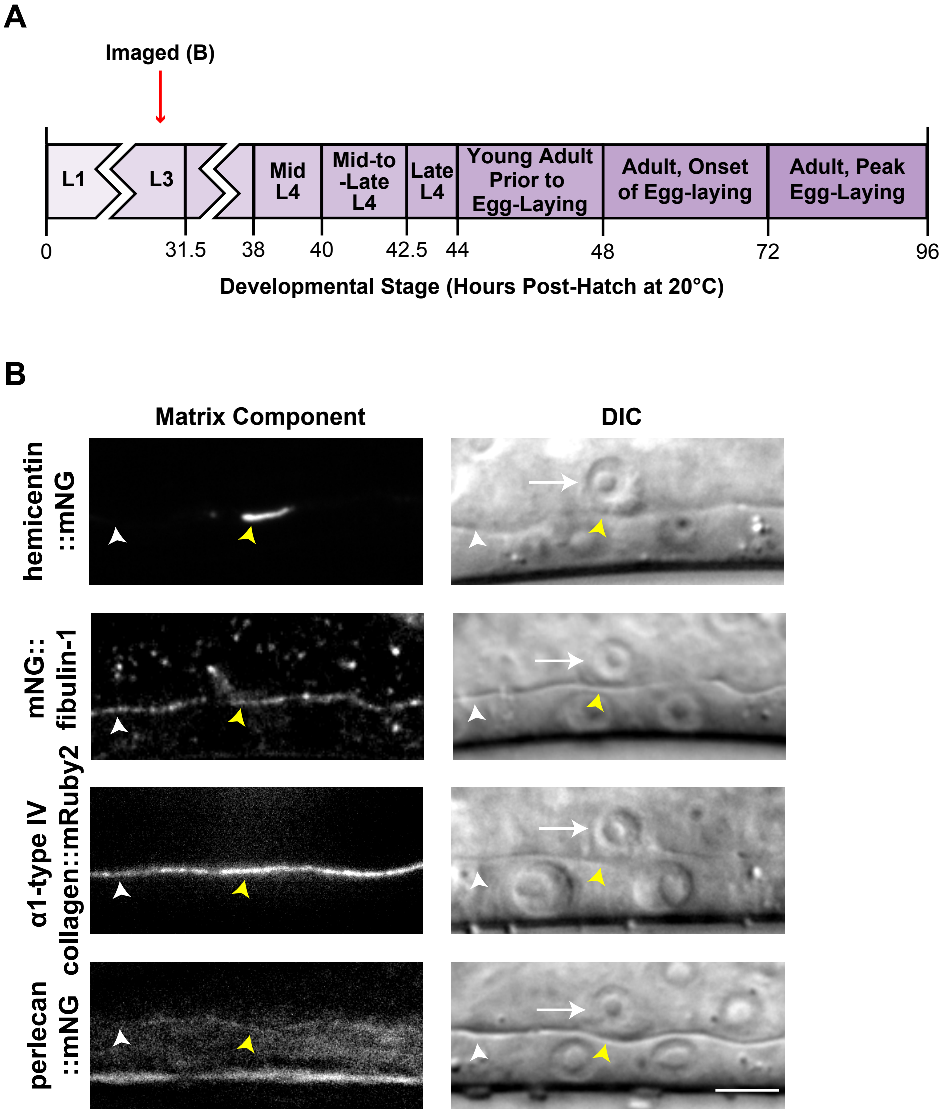
A
Imaged (B)
L1
L3
Mid L4
Mid-to -Late L4
Late L4
Young Adult Prior to Egg-Laying
Adult, Onset of Egg-laying
Adult, Peak Egg-Laying
0
31.5
38
40
42.5
44
48
72
96
Developmental Stage (Hours Post-Hatch at 20°C)
B
Matrix Component
DIC
hemicentin ::mNG
mNG:: fibulin-1
α1-type IV collagen::mRuby2
perlecan ::mNG

### Figure S2

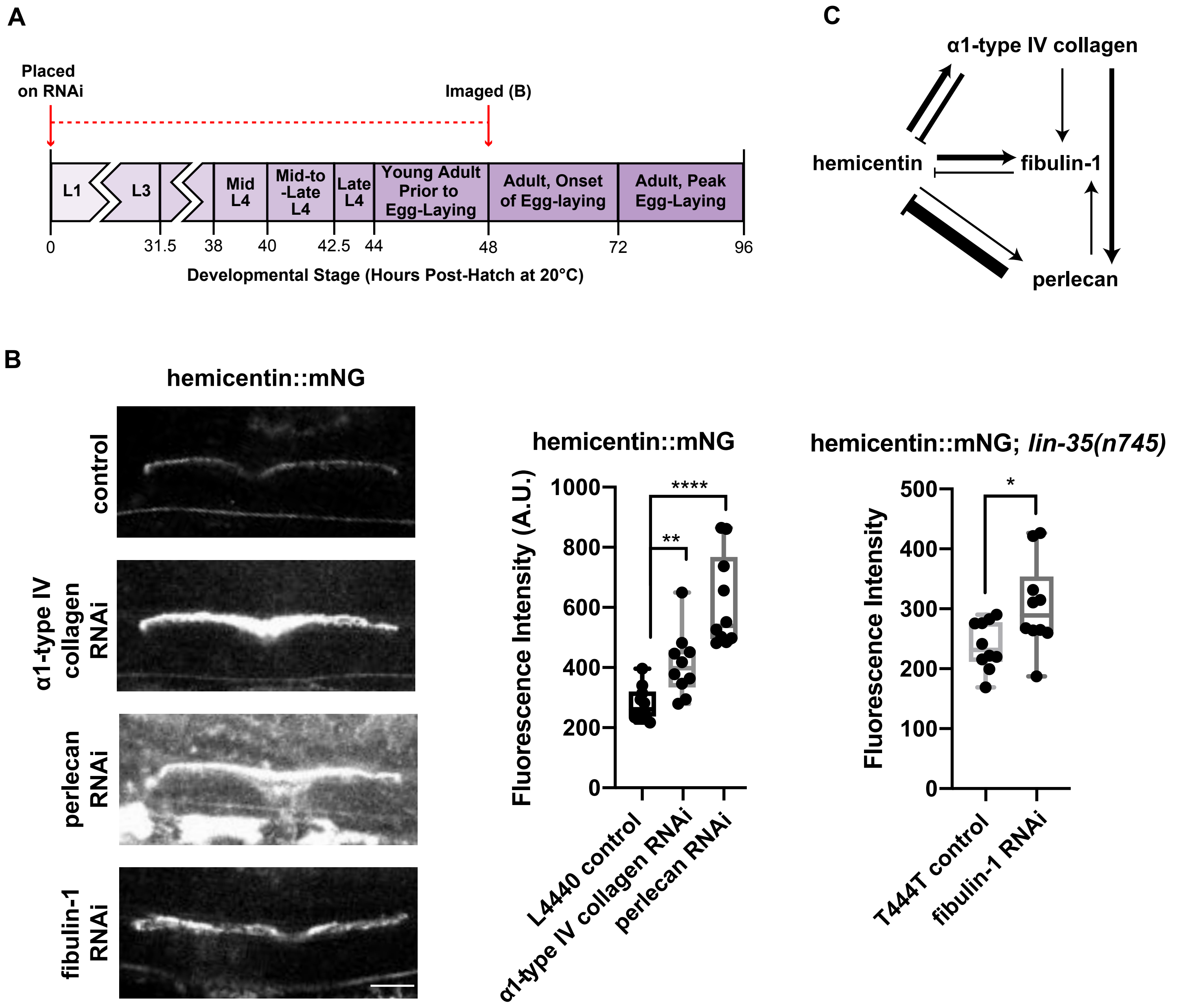
A
Placed on RNAi
Imaged (B)
L1
L3
Mid L4
Mid-to -Late L4
Late L4
Young Adult Prior to Egg-Laying
Adult, Onset of Egg-laying
Adult, Peak Egg-Laying
0
31.5
38
40
42.5
44
48
72
96
Developmental Stage (Hours Post-Hatch at 20°C)
C
α1-type IV collagen
hemicentin
fibulin-1
perlecan
B
hemicentin::mNG
control
α1-type IV collagen RNAi
perlecan RNAi
fibulin-1 RNAi
hemicentin::mNG
Fluorescence Intensity (A.U.)
1000
800
600
400
200
0
****
**
L4440 control
α1-type IV collagen RNAi
perlecan RNAi
hemicentin::mNG; lin-35(n745)
Fluorescence Intensity
500
400
300
200
100
0
*
T444T control
fibulin-1 RNAi

### Figure S3

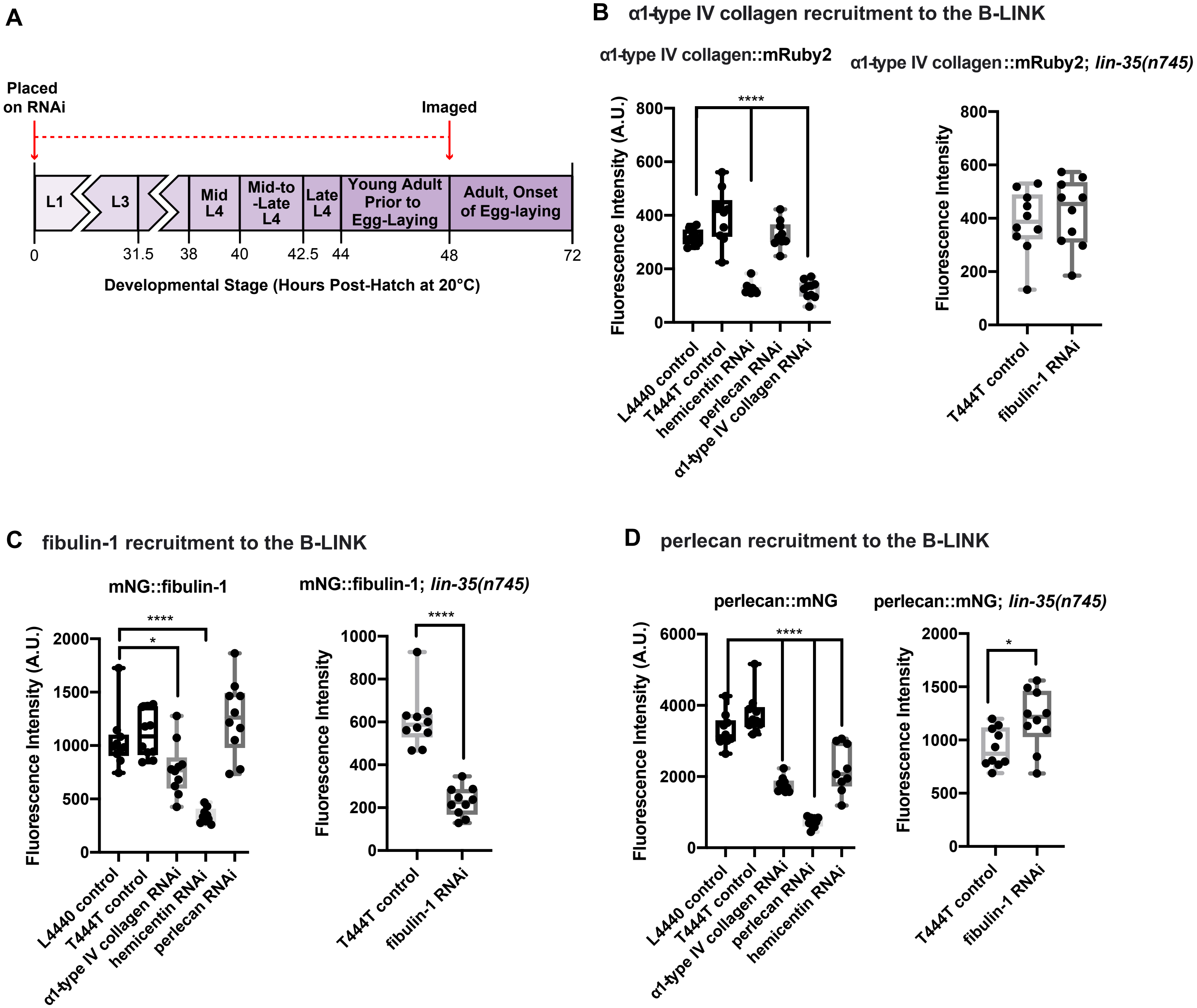
A
Placed on RNAi
Imaged
L1
L3
Mid L4
Mid-to -Late L4
Late L4
Young Adult Prior to Egg-Laying
Adult, Onset of Egg-laying
0
31.5
38
40
42.5
44
48
72
Developmental Stage (Hours Post-Hatch at 20°C)
B α1-type IV collagen recruitment to the B-LINK
α1-type IV collagen::mRuby2
α1-type IV collagen::mRuby2; lin-35(n745)
Fluorescence Intensity (A.U.)
Fluorescence Intensity
****
L4440 control
T444T control
hemicentin RNAi
perlecan RNAi
α1-type IV collagen RNAi
T444T control
fibulin-1 RNAi
C fibulin-1 recruitment to the B-LINK
mNG::fibulin-1
mNG::fibulin-1; lin-35(n745)
Fluorescence Intensity (A.U.)
Fluorescence Intensity
****
*
****
L4440 control
T444T control
α1-type IV collagen RNAi
hemicentin RNAi
perlecan RNAi
T444T control
fibulin-1 RNAi
D perlecan recruitment to the B-LINK
perlecan::mNG
perlecan::mNG; lin-35(n745)
Fluorescence Intensity (A.U.)
Fluorescence Intensity
****
*
L4440 control
T444T control
α1-type IV collagen RNAi
perlecan RNAi
hemicentin RNAi
T444T control
fibulin-1 RNAi

#### Figure S4

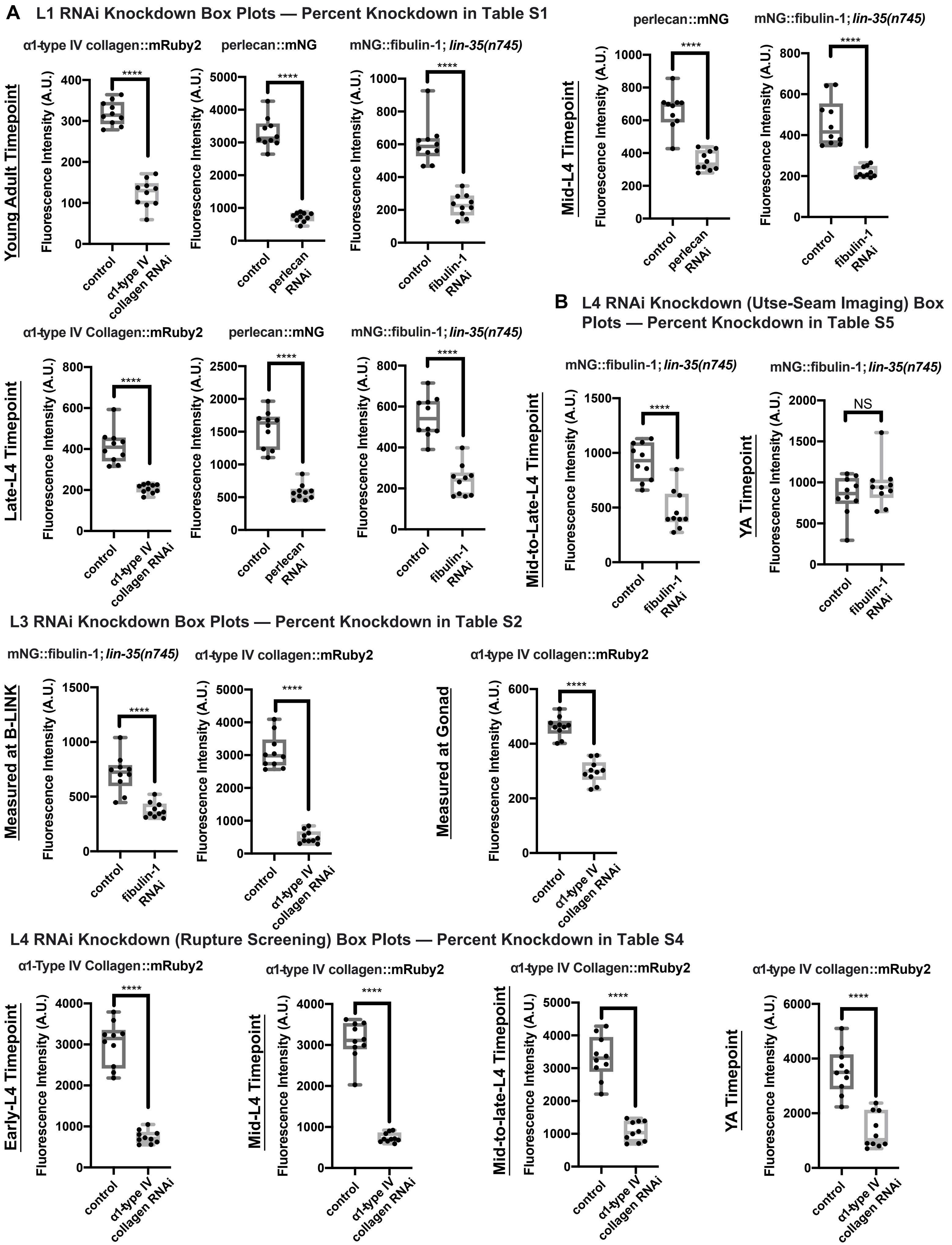

A
L1 RNAi Knockdown Box Plots — Percent Knockdown in Table S1
Young Adult Timepoint
α1-type IV collagen::mRuby2
perlecan::mNG
mNG::fibulin-1; lin-35(n745)
Fluorescence Intensity (A.U.)
****
control
α1-type IV collagen RNAi
perlecan RNAi
fibulin-1 RNAi
Mid-L4 Timepoint
perlecan::mNG
mNG::fibulin-1; lin-35(n745)
Late-L4 Timepoint
α1-type IV Collagen::mRuby2
perlecan::mNG
mNG::fibulin-1; lin-35(n745)
B
L4 RNAi Knockdown (Utse-Seam Imaging) Box Plots — Percent Knockdown in Table S5
Mid-to-Late-L4 Timepoint
YA Timepoint
mNG::fibulin-1; lin-35(n745)
NS
C
L3 RNAi Knockdown Box Plots — Percent Knockdown in Table S2
Measured at B-LINK
Measured at Gonad
mNG::fibulin-1; lin-35(n745)
α1-type IV collagen::mRuby2
D
L4 RNAi Knockdown (Rupture Screening) Box Plots — Percent Knockdown in Table S4
Early-L4 Timepoint
Mid-L4 Timepoint
Mid-to-late-L4 Timepoint
YA Timepoint
α1-Type IV Collagen::mRuby2
α1-type IV collagen::mRuby2
α1-type IV Collagen::mRuby2
α1-type IV collagen::mRuby2

Table S1. **Rupture screening of basement membrane matrix components**

| Gene[a] | Genotype/Treatment | % Ruptured[b] | *n*[c] | p-value[d] |
|---|---|---|---|---|
| **Mutant** | | | | |
| control | *N2* | 0% | 222 | NA |
| agrin | *agr-1(oxTi4)* | 0% | 76 | NS |
| nidogen | *nid-1(cg119)* | 0% | 93 | NS |
| type XVIII collagen | *cle-1(cg120)* | 0% | 58 | NS |
| perlecan | *unc-52(e998)* | 0% | 21 | NS |
| fibulin-1 | *fbl-1(hd42)* | 20% | 20 | 0.0001 |
| **L1 RNAi** | | | | |
| control | *L4440 (RNAi)* | 3% | 151 | NA |
| agrin | *agr-1(RNAi)* | 0% | 206 | NS |
| nidogen | *nid-1(RNAi)* | 0% | 200 | NS |
| spondin | *spon-1(RNAi)* | 0% | 241 | NS |
| type XVIII collagen | *cle-1(RNAi)* | 0% | 187 | NS |
| perlecan | *unc-52(RNAi)* | 0% | 198 | NS |
| fibulin-1 | *fbl-1(RNAi)* | 17% | 169 | 0.001 |
| papilin | *mig-6 (RNAi)* | 39% | 158 | 0.0001 |
| γ-laminin | *lam-2 (RNAi)* | 38% | 149 | 0.0001 |
| α1-type IV collagen | *emb-9 (RNAi)* | 84% | 147 | 0.0001 |
| hemicentin | *him-4 (RNAi)* | 93% | 152 | 0.0001 |
| **L3 RNAi** | | | | |
| control | *L4440 (RNAi)* | 2% | 66 | NA |
| hemicentin | *him-4 (RNAi)* | 97% | 34 | 0.0001 |
| α1-type IV collagen | *emb-9 (RNAi)* | 97% | 146 | 0.0001 |
| papilin | *mig-6 (RNAi)* | 2% | 85 | NS |
| γ-laminin | *lam-2 (RNAi)* | 3% | 130 | NS |
| fibulin-1 | *fbl-1(RNAi)* | 9% | 188 | 0.05 |
| perlecan | *unc-52(RNAi)* | 0% | 133 | NS |

[a] All RNAi was completed using a whole body RNAi sensitized strain (γ-laminin::mNG; rrf-3) with the exception of fibulin RNAi, which was completed in a strain with a null mutation in *lin-35* [*lin-35(n745)*], which improved RNAi efficiency.

[b] Ruptures were visually scored at least once every 24 hours for 120 hours post plating (See Methods).

[c] Number of animals scored per condition.

[d] P-values were calculated used Fisher's Exact Tests. Conditions were compared to corresponding empty vector/N2 controls. NS, not significant. NA, not applicable.

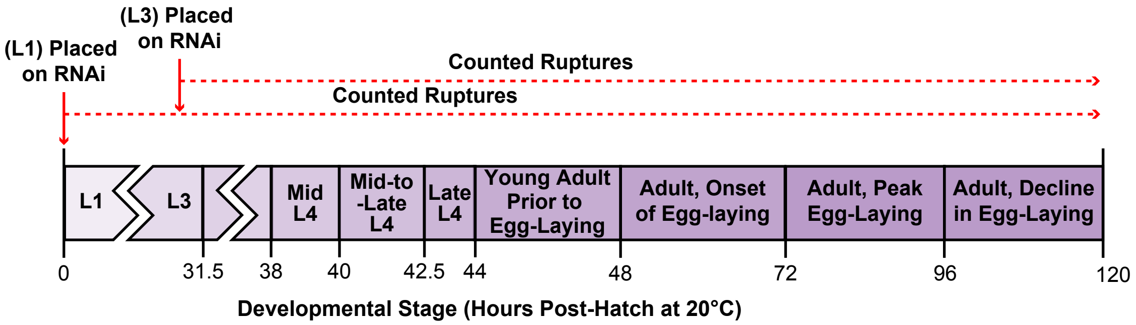

Table S2. **L1 RNAi knockdown efficiencies**

| Gene[a] | Genotype/Treatment | Time on RNAi | % Knockdown[b] | *n*[c] |
|---|---|---|---|---|
| **Imaged at Mid-L4** | | | | |
| α1-type IV collagen | *emb-9 (RNAi)* | 39 h | NA | NA |
| hemicentin | *him-4 (RNAi)* | 39 h | 100% | 10 |
| perlecan | *unc-52 (RNAi)* | 39 h | 55% | 10 |
| fibulin-1 | *fbl-1 (RNAi)* | 39 h | 67% | 10 |
| **Imaged at Late-L4** | | | | |
| α1-type IV collagen | *emb-9 (RNAi)* | 43 h | 66% | 10 |
| hemicentin | *him-4 (RNAi)* | 43 h | 100% | 10 |
| perlecan | *unc-52 (RNAi)* | 43 h | 67% | 10 |
| fibulin-1 | *fbl-1 (RNAi)* | 43 h | 69% | 10 |
| **Imaged at YA** | | | | |
| α1-type IV collagen | *emb-9 (RNAi)* | 48 h | 62% | 10 |
| hemicentin | *him-4 (RNAi)* | 48 h | 100% | 10 |
| perlecan | *unc-52 (RNAi)* | 48 h | 78% | 10 |
| fibulin-1 | *fbl-1 (RNAi)* | 48 h | 69% | 10 |

[a] All RNAi was completed using endogenous fluorophore tagged lines (α1-type IV collagen::mRuby2, hemicentin::mNG, perlecan::mNG) with the exception of fibulin RNAi, which was completed in a strain with a null mutation in *lin-35* [mNG::fibulin; *lin-35(n745)*], which improved RNAi efficiency. YA, young adult.

[b] Knockdown was calculated by taking the mean fluorescence intensity of a 3-pixel wide line drawn through the left side of the B-LINK (See Methods). Box plots with knockdown efficency data are shown in Fig. S4. NA, not applicable (component was not present in controls or RNAi conditions at given timepoint). 100% indicates signal was undetectable after RNAi knockdown.

[c] Number of animals scored per condition. NA, not applicable

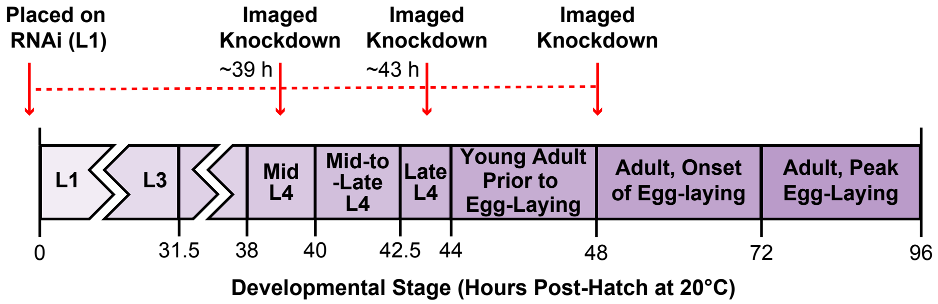

Table S3. **L3 RNAi knockdown efficiencies**

| Gene[a] | Genotype/Treatment | Time on RNAi | % Knockdown[b] | *n*[c] |
|---|---|---|---|---|
| **B-LINK** | | | | |
| α1-type IV collagen | *emb-9 (RNAi)* | 91 h | 87% | 10 |
| hemicentin | *him-4 (RNAi)* | 91 h | 100% | 10 |
| fibulin-1 | *fbl-1(RNAi)* | 91 h | 53% | 10 |
| **Gonad** | | | | |
| α1-type IV collagen | *emb-9 (RNAi)* | 91 h | 45% | 10 |

[a] All RNAi was completed using endogenous fluorophore tagged lines (α1-type IV collagen::mRuby2, hemicentin::mNG, perlecan::mNG) with the exception of fibulin RNAi, which was completed in a strain with a null mutation in *lin-35* [mNG::fibulin; *lin-35(n745)*], which improved RNAi efficiency.

[b] Knockdown was calculated by taking the mean fluorescence intensity of a 3-pixel wide line drawn through the left side of the B-LINK (See Methods). Box plots with knockdown efficency data are shown in Fig. S4. 100% indicates signal was undetectable after RNAi knockdown.

[c] Number of animals scored per condition.

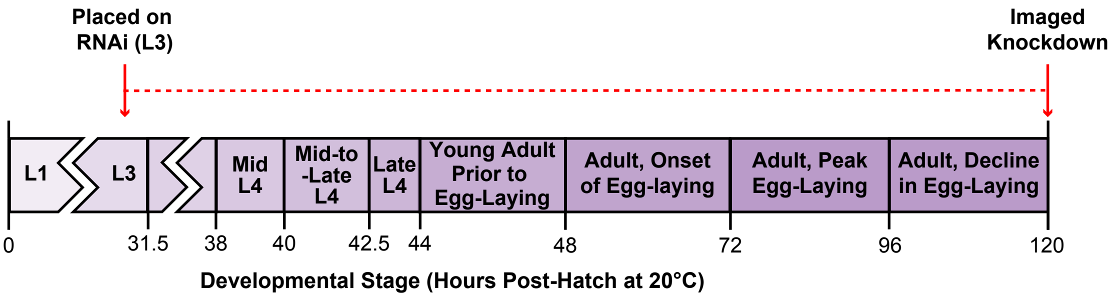

Table S4. **Rupture screening of basement membrane matrix components**

| Gene[a] | Genotype/Treatment | % Ruptured[b] | *n*[c] | p-value[d] |
|---|---|---|---|---|
| **L1 Uterine Specific RNAi*** | | | | |
| control | *L4440 (RNAi)* | 1% | 109 | NA |
| hemicentin | *him-4 (RNAi)* | 80% | 79 | 0.0001 |
| α1-type IV collagen | *emb-9 (RNAi)* | 1% | 85 | NS |
| papilin | *mig-6 (RNAi)* | 3% | 342 | NS |
| γ-laminin | *lam-2 (RNAi)* | 2% | 86 | NS |
| fibulin-1 | *fbl-1(RNAi)* | 1% | 92 | NS |
| perlecan | *unc-52(RNAi)* | 1% | 116 | NS |
| **L1 Hypodermal Specific RNAi*** | | | | |
| control | *L4440 (RNAi)* | 0% | 164 | NA |
| hemicentin | *him-4 (RNAi)* | 0% | 155 | NS |
| α1-type IV collagen | *emb-9 (RNAi)* | 0% | 178 | NS |
| papilin | *mig-6 (RNAi)* | 0% | 172 | NS |
| perlecan | *unc-52(RNAi)* | 0% | 160 | NS |

[a] Uterine specific RNAi was completed using a uterine specific RNAi strain (see Table S7). Hypodermal specific RNAi was completed using a hypodermal specific RNAi strain (see Table S7).

[b] Ruptures were visually scored at least once every 24 hours for 120 hours post plating (See Methods).

[c] Number of animals scored per condition.

[d] P-values were calculated used Fisher's Exact Tests. Conditions were compared to corresponding empty vector controls. NS, not significant. NA, not applicable.

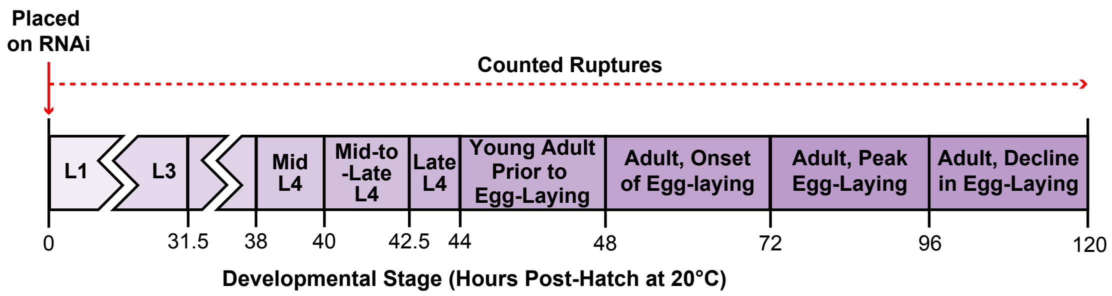

Table S5. **L4 RNAi knockdown efficiencies, rupture experiments**

| Gene[a] | Genotype/Treatment | Time on RNAi | % Knockdown[b] | *n*[c] |
|---|---|---|---|---|
| **<u>Early-L4 RNAi</u> (35 h)** | | | | |
| α1-type IV collagen | *emb-9 (RNAi)* | 85 h | 81% | 10 |
| hemicentin | *him-4 (RNAi)* | 85 h | 100% | 10 |
| **<u>Mid-L4 RNAi</u> (37.5 h)** | | | | |
| α1-type IV collagen | *emb-9 (RNAi)* | 82.5 h | 79% | 10 |
| hemicentin | *him-4 (RNAi)* | 82.5 h | 100% | 10 |
| **<u>Mid-to-Late-L4 RNAi</u> (40 h)** | | | | |
| α1-type IV collagen | *emb-9 (RNAi)* | 80 h | 70% | 10 |
| hemicentin | *him-4 (RNAi)* | 80 h | 100% | 10 |
| **<u>Young Adult RNAi</u> (48 h)** | | | | |
| α1-type IV collagen | *emb-9 (RNAi)* | 72 h | 63% | 10 |
| hemicentin | *him-4 (RNAi)* | 72 h | 100% | 10 |

[a] All RNAi was completed using endogenous fluorophore tagged lines (α1-type IV collagen::mRuby2, hemicentin::mNG, perlecan::mNG) with the exception of fibulin RNAi, which was completed in a strain with a null mutation in *lin-35* [mNG::fibulin; *lin-35(n745)*], which improved RNAi efficiency.

[b] Knockdown was calculated by taking the mean fluorescence intensity of a 3-pixel wide line drawn through the left side of the B-LINK (See Methods). Box plots with knockdown efficency data are shown in Fig. S4. 100% indicates signal was undetectable after RNAi knockdown.

[c] Number of animals scored per condition.

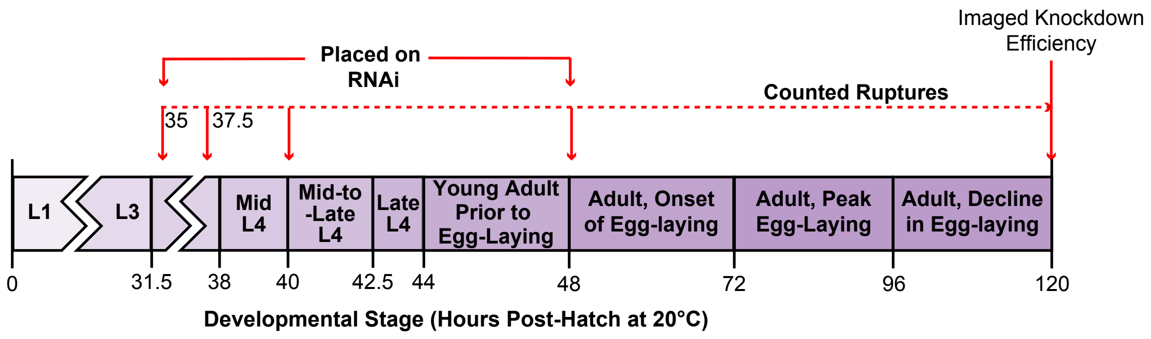

Table S6. **L4 RNAi knockdown efficiencies, utse-seam imaging**

| Gene[a] | Genotype/Treatment | Time on RNAi | % Knockdown[b] | *n*[c] |
|---|---|---|---|---|
| **<u>Mid-to-Late-L4 RNAi</u> (40 h)** | | | | |
| fibulin-1 | *fbl-1 (RNAi)* | 32 h | 52% | 10 |
| **<u>Young Adult RNAi</u> (48 h)** | | | | |
| hemicentin | *him-4 (RNAi)* | 24 h | 100% | 10 |
| fibulin-1 | *fbl-1 (RNAi)* | 24 h | 1% | 10 |

[a] hemicentin RNAi was completed using an endogenous fluororphore tagged line (hemicentin::mNG) fibulin RNAi was completed in a strain with a null mutation in *lin-35* [mNG::fibulin; *lin-35(n745)*], which improved RNAi efficiency.

[b] Knockdown was calculated by taking the mean fluorescence intensity of a 3-pixel wide line drawn through the left side of the B-LINK (See Methods). Box plots with knockdown efficency data are shown in Fig. S3. 100% indicates signal was undetectable after RNAi knockdown.

[c] Number of animals scored per condition.

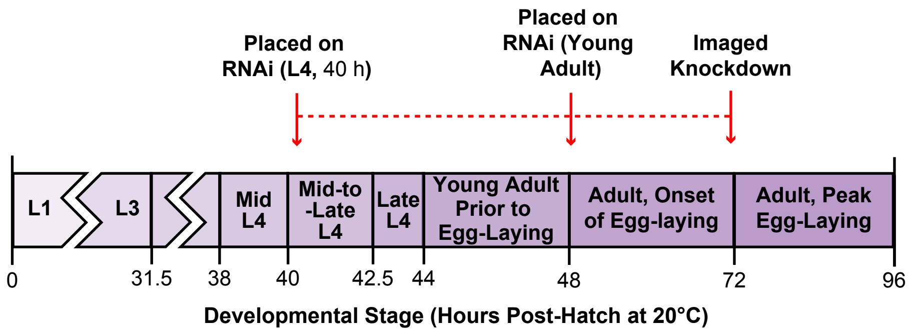

Table S7: **Strain information**

| Strain | Genotype[a] | Reference |
|---|---|---|
| NK2617 | ***qyIs23** [cdh-3p::mCh::PH]* II*;* ***lqIs80** [SCMp::GFP::CAAX]* IV | This study |
| NK2422 | ***qy33** [him-4::mNG]* X | Keeley et al., 2020 |
| NK2324 | ***qy23** [ina-1::mNG]* III | Jayadev et al., 2019 |
| NK1593 | ***qy9** (vab-10a::GFP* + *unc-119(+))* | Morrissey et al., 2014 |
| NK2583 | ***qy80** [unc-52::mNG]* II | Keeley et al., 2020 |
| NK2579 | ***qy62** [mNG::fbl-1]* IV | Keeley et al., 2020 |
| NK2326 | ***qy24** [emb-9::mNG]* III | Keeley et al., 2020 |
| NK2556 | ***qy72** [mig-6S::mNG]/*+ V | Keeley et al., 2020 |
| NK2443 | ***qy38** [nig-1::mNG]* V | Keeley et al., 2020 |
| NK2322 | ***qy22** [cle-1::mNG]* I | Keeley et al., 2020 |
| NK2557 | ***qy73** [mig-6L::mNG]* V | Keeley et al., 2020 |
| NK2580 | ***qy30** [spon-1::mNG]* II | Keeley et al., 2020 |
| NK2335 | ***qy20** [lam-2::mNG]* X | Jayadev et al., 2019 |
| KG1271 | *lite-1(ce314)* X*;* ***ceIs37** [myo-3p::lite-1* + *myo-3p::GFP]* X | Edwards et al., 2008 |
| NK2585 | ***qy83** [emb9::mRuby2]* III | Jayadev et al., 2021 |
| NK2689 | *lin-35(n745)* I*; qyIs23 II; lqIs80* IV | This study |
| NK2645 | *lin-35(n745)* I*; qy33* X | This study |
| NK2651 | *lin-35(n745)* I*; qy83* III | This study |
| NK2644 | *lin-35(n745)* I*; qy62* IV | This study |
| NK2643 | *lin-35(n745)* I*; qy80* II | This study |
| N2 | Wild-type (ancestral) | --- |
| EG1770 | *agr-1(oxTi4)* II | Hrus et al., 2007 |
| CH119 | *nid-1(cg119)* V | Kang et al., 2000 |
| CH120 | *cle-1(cg120)* I | Ackley et al., 2001 |
| CB998 | *unc-52(e998)* II | Rogalski et al., 1994 |
| NK1905 | *fbl-1(hd43)/nT1* IV, (IV,V) | Muriel et al., 2005 |
| NK2334 | *rrf-3(pk1426)* II*; qy20* X | Jayadev et al., 2019 |
| NK1316 | *rrf-3(pk1426)* II*;* ***qyIs102** [fos-1ap::RDE-1, myo-2p::GFP];* ***qyIs10** [lam-1p::lam-1::GFP]* IV*; rde-1 (ne219)* V*;* ***qyIs24** [cdh-3p::mCherry::PLCdPH]* | Morrissey et al., 2014 |
| NR222 | *rde-1(ne219)* V*;* ***kzIs9**[pKK1260(lin-26p::nls::gfp), pKK1253(lin-26p::rde-1), pRF4(rol-6 marker)]* | Qadota et al., 2007 |
| NK970 | ***rhIs23** [him-4::GFP] X;* ***qyIs50*** [cdh-3::mCh::moeABD] II; rol-6(su1009) | Morrissey et al., 2014 |

[a] When a transgene is listed in the table for the first time, it is bolded and the entire genotype is displayed.
